# Intrinsic and growth-mediated cell and matrix specialization during meniscus tissue assembly

**DOI:** 10.1101/2021.02.20.432114

**Authors:** Tonia K. Tsinman, Xi Jiang, Lin Han, Eiki Koyama, Robert L. Mauck, Nathaniel A. Dyment

## Abstract

The incredible mechanical strength and durability of mature fibrous tissues and their extremely limited turnover and regenerative capacity underscores the importance of proper matrix assembly during early postnatal growth. In tissues with composite extracellular matrix (ECM) structures, such as the adult knee meniscus, fibrous (Collagen-I rich) and cartilaginous (Collagen-II, proteoglycan-rich) matrix components are regionally segregated to the outer and inner portions of the tissue. While this spatial variation in composition is appreciated to be functionally important for resisting complex mechanical loads associated with gait, the establishment of these specialized zones is poorly understood. To address this issue, the following study tracked the growth of the murine meniscus from its embryonic formation through its first month of growth, encompassing the critical time-window during which animals begin to ambulate and weight bear. Using histological analysis, region specific high-throughput qPCR, and Col-1 and Col-2 fluorescent reporter mice, we found that matrix and cellular features defining specific tissue zones were already present at birth, before continuous weight-bearing had occurred. These differences were further refined with postnatal growth and maturation, resulting in specialization of mature tissue regions. Taken together, this work establishes a detailed timeline of the concurrent spatiotemporal changes that occur at both the cellular and matrix level throughout meniscus maturation. The findings of this study provide a framework for investigating the reciprocal feedback between cells and their evolving microenvironments during assembly of a mechanically robust fibrocartilage tissue, thus providing insight into mechanisms of tissue degeneration and effective regenerative strategies.

## INTRODUCTION

It is well established that proper function of mature tissues is dependent on their cellular and molecular composition and organization. Classic examples, such as the intestinal villi, pancreatic islets, hepatic lobules, and mammary acini, all demonstrate how the precise assembly of specialized cells into a supporting extracellular matrix (ECM) enables the execution of unique physiologic tasks such as nutrient absorption, hormone secretion, and metabolism (Malarkey *et al*., 2005; Kass *et al*., 2007; Clevers, 2013; Llacua, Faas and de Vos, 2018). Of these specialized functions, one requiring an exquisitely engineered tissue, is locomotion. Articular cartilage, tendons, ligaments, and the menisci of the knee joint, repeatedly experience mechanical loads up to several times an animal’s body weight. In their mature state, these tissues rely on the robust mechanical properties of their established ECMs; with each structure finely tuned to withstand the specialized forces needed for native function. The matrix within ligaments and tendons, for instance, contains linearly aligned, thick fibrous Collagen-I (Col-1) bundles that effectively resist uniaxial tensile forces associated with muscle-to-bone force transmission (Benjamin, Kaiser and Milz, 2008; Screen *et al*., 2015). Conversely, articular cartilage (e.g., the lining of the femur and tibia) has a relatively isotropic matrix organization comprised predominantly of Collagen-II (Col-2) fibrils. This Col-2 rich composition is complemented with an abundance of negatively charged proteoglycans (PGs), which engenders a high water content and enables the resistance to large compressive forces generated within the knee during each gait cycle (Sophia Fox, Bedi and Rodeo, 2009; Roughley and Mort, 2014; Wilusz, Sanchez-Adams and Guilak, 2014).

While cartilage (predominantly compressive) and ligament (predominantly tensile) represent the two extremes in mechanical specialization of musculoskeletal ECM composition, other tissues reside on this structure-function spectrum. In particular, the knee meniscus is a semilunar wedge-shaped structure between the femoral condyles and tibial plateau that has a composite ECM structure, containing both fibrous and cartilaginous components (Valiyaveettil, Mort and McDevitt, 2005; Chevrier *et al*., 2009). As the meniscus serves as a central load transmitting tissue during knee joint motion, the regional variation in ECM complements the need to withstand both compression from the adjacent femoral and tibial surfaces, as well as the tension within circumferentially oriented collagen fibers that resist extrusion of the tissue from the joint space (Fox, Bedi and Rodeo, 2012; Andrews *et al*., 2017). The localization of the matrix components is precisely defined to meet these mechanical demands, with the outer portion of the wedge containing circumferentially aligned Col-1-rich fibers while the inner portion of the tissue contains matrix with more Col-2 with higher PG-rich content (Kambic and McDevitt, 2005; Killian *et al*., 2010; Makris, Hadidi and Athanasiou, 2011; Vanderploeg *et al*., 2012; Andrews *et al*., 2013). While the functional importance of these inner-to-outer matrix differences is well appreciated, how such a composite ECM structure is assembled by the resident cell population and when distinctions between these regions first arise remains unclear. In fact, a breakdown of this regionalization, whereby PG-rich matrix appears in outer regions normally occupied by the circumferential fibers is associated with aging and degeneration and a loss in mechanical integrity (Hellio Le Graverand *et al*., 2001; M. P. Hellio Le Graverand *et al*., 2001b, 2001a; Pauli *et al*., 2011; Han *et al*., 2016; Kwok *et al*., 2016). Thus, determining how a regionally distinct ECM is assembled may not only shed light on principles driving the formation of complex composite ECM structures, but also may illuminate operative mechanisms of meniscus pathogenesis and guide therapeutic intervention.

Though the mature meniscus has a relatively low cell content, the assembly, organization, and maintenance of the complex ECM network is a cell-mediated process. Yet, whether the divergent matrix synthesis in the inner and outer region is due to an adaptation of local cells to differential biophysical cues or if these regions represent distinct subpopulations of meniscus cells that are pre-fated towards a more chondrogenic or a fibrogenic phenotype is not yet known. Interestingly, embryonic and neonatal menisci of large animals are mostly fibrous, and PG enrichment in the inner ECM region occurs gradually during postnatal growth (Melrose *et al*., 2005; Smith, Shu and Melrose, 2010; Ionescu *et al*., 2011; Di Giancamillo *et al*., 2014). Because the early period of postnatal tissue growth coincides with the time at which animals become increasingly motile, and PGs are crucial for effective dissipation of compressive loads (as in articular cartilage), a prevailing theory is that inner region synthesis of cartilage-like ECM is a response of the local inner meniscus cells to compressive forces. Indeed, cells within inner and outer meniscus regions show varied morphology and expression signatures, and growing evidence suggests that the cellular microenvironment – including its composition, topology, stiffness, and subjection to extrinsic forces – is a significant contributor to directing and maintaining cellular phenotype (Hellio Le Graverand *et al*., 2001; Son and Levenston, 2012; McNulty and Guilak, 2015; Kumar, Placone and Engler, 2017). Contrary to this hypothesis, however, lineage tracing studies using mouse models have suggested that cells within the meniscus are of mixed origin (Hyde, Boot-Handford and Wallis, 2008), indicating that spatial differences in ECM synthesis by cells may be specified by cell patterning during tissue formation. While the murine meniscus model has proven to be a powerful tool for studying molecular contributions to meniscus formation and defining the critical timepoints during development, concurrent assessment of regional changes in cellular phenotype and matrix properties has not been reported (Hyde, Boot-Handford and Wallis, 2008; Pazin *et al*., 2014; Shwartz *et al*., 2016; Gamer *et al*., 2017, 2018; Gamer, Xiang and Rosen, 2017).

To that end, we herein define the spatiotemporal specialization of the developing murine meniscus ECM in parallel with the time-evolving molecular profiles of the resident cell subpopulations in the late embryonic and early post-natal time frame. We utilize a *Col1*-YFP/*Col2*-CFP reporter mouse model to identify patterns of expression of the two major collagen types within the meniscus at single cell resolution and analyze regional meniscus transcriptional profiles and matrix properties relative to adjacent ligamentous and cartilaginous tissues within the knee joint. Our findings show that regional distinctions in both matrix and cellular fate are present at birth, prior to continuous weight-bearing, and that both cells and matrix further specialize over time. These findings suggest that regional specialization occurs early in meniscus embryonic formation and highlights the dramatic changes in tissue properties that occur over time spans as short as one week. Taken together, this work establishes a detailed timeline of the concurrent spatiotemporal changes that occur at both the cellular and matrix level throughout meniscus maturation, providing a foundation for elucidating the regulatory feedback loop between the mechanical and structural features of the ECM microenvironment and endogenous cellular phenotypes.

## RESULTS

### The murine meniscus exhibits biphasic growth patterns during the early postnatal period

We began our analysis of tissue maturation by quantifying the growth characteristics of the body region of the meniscus from embryonic formation (E17.5) to adulthood (4 months). Our specific focus was on the first 4 weeks of postnatal growth (P0-P28), the time period over which the cross-sectional area of the meniscus increases to ∼85% of its adult state. Because of the unique semilunar, wedge-shaped geometry of the tissue, we performed our analysis in the coronal plane, where the meniscus cross-section could be observed relative to the adjacent tissues within the joint space (**Fig. 1a**). Importantly, this resulted in the inner and outer regions of the tissue being visualized within a single sectioning plane (**Fig. 1a; Fig. S1a**). We noted a rapid increase in coronal cross-sectional area from E17.5 to P14 – especially in the outer half of the tissue – followed by a plateau in the growth rate (**Fig. 1b,c; Fig. S1b**), consistent with previous observations (Gamer, Xiang and Rosen, 2017). Notably, the expansion of the meniscus wedge was asymmetric, as the ratio of height to length decreased with time. This was mirrored by an increase in the tibial plateau coverage – indicating a lengthening of the tissue towards the midline of the joint (**Fig. 1d; Fig. S1c, d**), as occurs in humans (Clark and Ogden, 1983). Observed growth in both the inner and outer regions was largely not due to cellular proliferation, as evidenced by the low number of proliferating cells over this time period and the decrease in cellularity (**Fig. 1e,f; Fig. S1b**). With respect to growth kinetics, all measured parameters followed a similar trajectory, with the most significant changes detected between P0 and P14, followed by a plateau from P14 onwards. Together, these initial studies point to two distinct growth phases of the mouse meniscus, with the early postnatal stage (P0-P14) characterized by rapid shifts in tissue size, shape, and cellularity, and the later postnatal stage (P14-adulthood) exhibiting only modest changes in these parameters.

**Figure 1:**
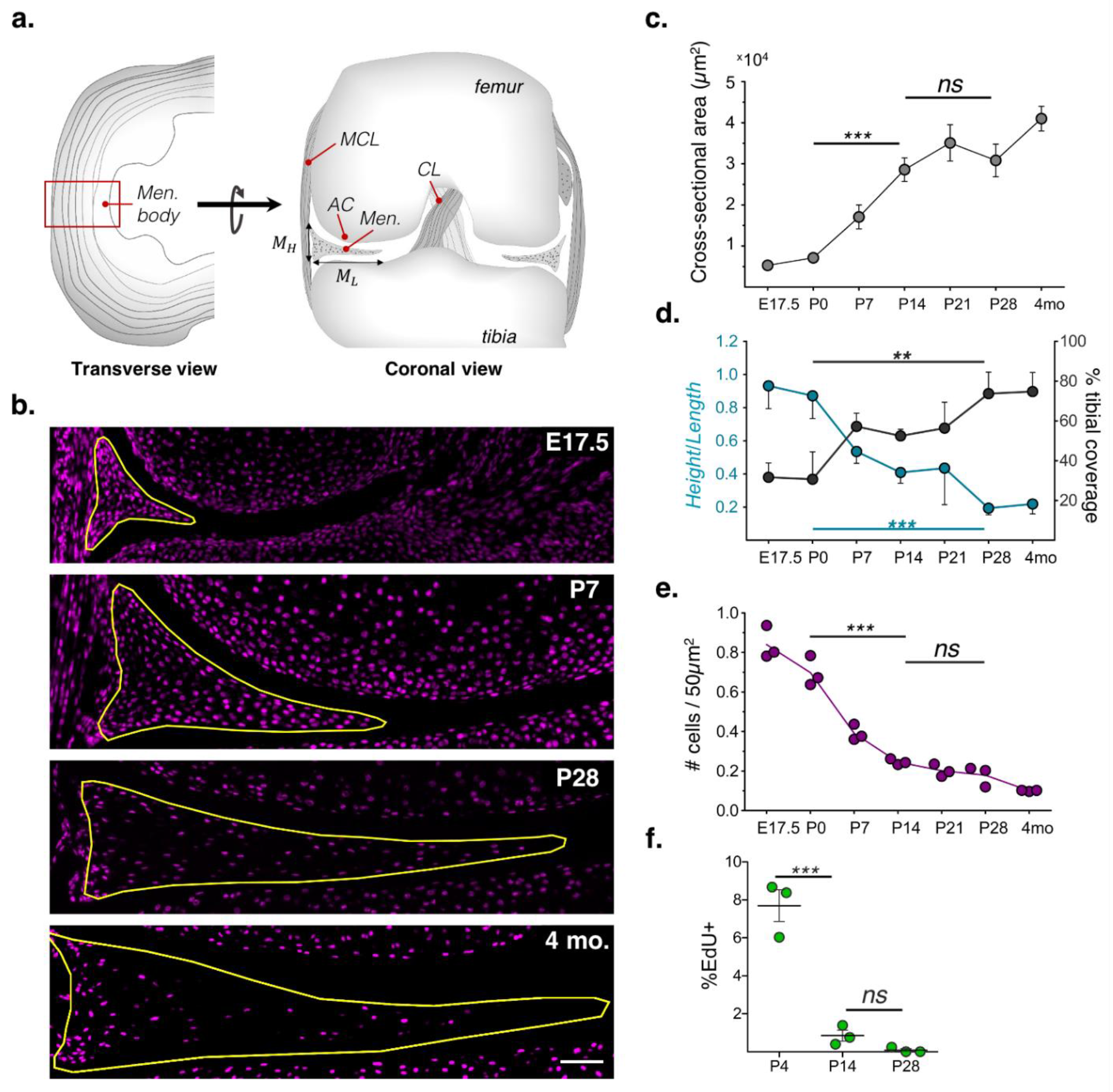
The murine meniscus undergoes dramatic growth and shape changes during the first month of postnatal growth. **a)** (left) Schematic of the transverse view of the meniscus tissue, demonstrating the plane in which the stereotypic semilunar structure, and circumferential collagen fibers, can be visualized. Red box indicates the meniscus ‘body’ region analyzed in this study. (right) Schematic of the coronal view of the knee joint, indicating the location of the meniscus relative to other important load bearing structures of the knee. AC: articular cartilage; MCL: medial collateral ligament; CL: cruciate ligament. M_L_, M_H_ length and height of the meniscus coronal cross-section, respectively. See figure supplement for full measurement annotation. **b)** Growth and shape change of the meniscus demonstrated through coronal-plane meniscus sections at specified timepoints, counterstained with DAPI (magenta). Yellow outline indicates meniscus cross-section. Scale bar: 100μm. **c,d)** Quantification of meniscus tissue growth and elongation into the joint space: **(c)** coronal cross sectional area, (**d)** ratio of meniscus height to its length (blue line, M_H_/M_L_) and percent of the medial tibial plateau covered by the meniscus(grey line, % tibial coverage). Mean ± sd shown. **e)** Number of cells per 50μm_2_ of the total cross sectional area (red line), with each biological replicate plotted. **f)** Percentage of cells per cross-section positive for EdU staining, indicating proliferative cells. n=3 biological replicates per timepoint. **: p<0.01; ***: p<0.0001, ns: not significant by one-way ANOVA with Tukey post-hoc.

### Region-specific meniscus matrix structure and composition is present at birth and evolves with maturation

The above data support that, consistent with other fibrous tissues, postnatal meniscus expansion occurs as a consequence of ECM accumulation rather than cell proliferation (Grinstein *et al*., 2019). To evaluate ECM content and structure during growth, we assessed proteoglycan distribution (via histological staining) and organized collagenous matrix (via both histological staining and second harmonic generation (SHG) multiphoton imaging). At birth (P0), proteoglycans were already enriched in the inner portion of the tissue, while collagen staining was strongest in the outer region (**Fig. 2a,b; Fig. 2 S1b**). SHG imaging at this early time point revealed that highly aligned circumferential fibers were evident at birth as well and were restricted to the outer region (**Fig. 2c; Movie 1**). Interestingly, while proteoglycan distribution remained similar at P7, fibrillar collagen rapidly accumulated in the inner region during the first week of growth (**Fig. 2c**). Over this same timeframe, cells in both regions became increasingly encased in fibrillar matrix, such that individual lacunae could be observed (**Movie 2; Fig. 2 S1e**). At later stages of maturation (P14-P28), a dense, circumferential fibrous matrix was well established throughout the meniscus body (**Fig. 2b; Fig. 2 S1c,d**). By P14, the proteoglycan-rich inner margin (seen at P0-P7) was less distinct. Rather, at this later stage, proteoglycans were most evident in the pericellular space (**Fig. 2a; Fig. 2 S1b**). This pericellular proteoglycan-rich matrix percolated outward with maturation, such that by P28, cells throughout the meniscus body showed dense local proteoglycan staining (**Fig. 2a**). As mouse pups become increasingly motile after P7, the presence of proteoglycans in the outer meniscus at later stages of maturation may indeed indicate a shift in cellular matrix synthesis in response to larger portions of the meniscus undergoing compressive loading. Additionally, given the increased abundance of dense fibrillar matrix with growth, the localization of the proteoglycan staining around the cells may be the result of the collagen matrix restricting newly synthesized ECM components to the pericellular space. Taken together, these data indicated that by birth, cells of the inner and outer meniscus already reside in microenvironments of distinct structural and molecular composition, that continue to evolve as the tissue grows and accrues a dense fibrous matrix topology.

**Figure 2:**
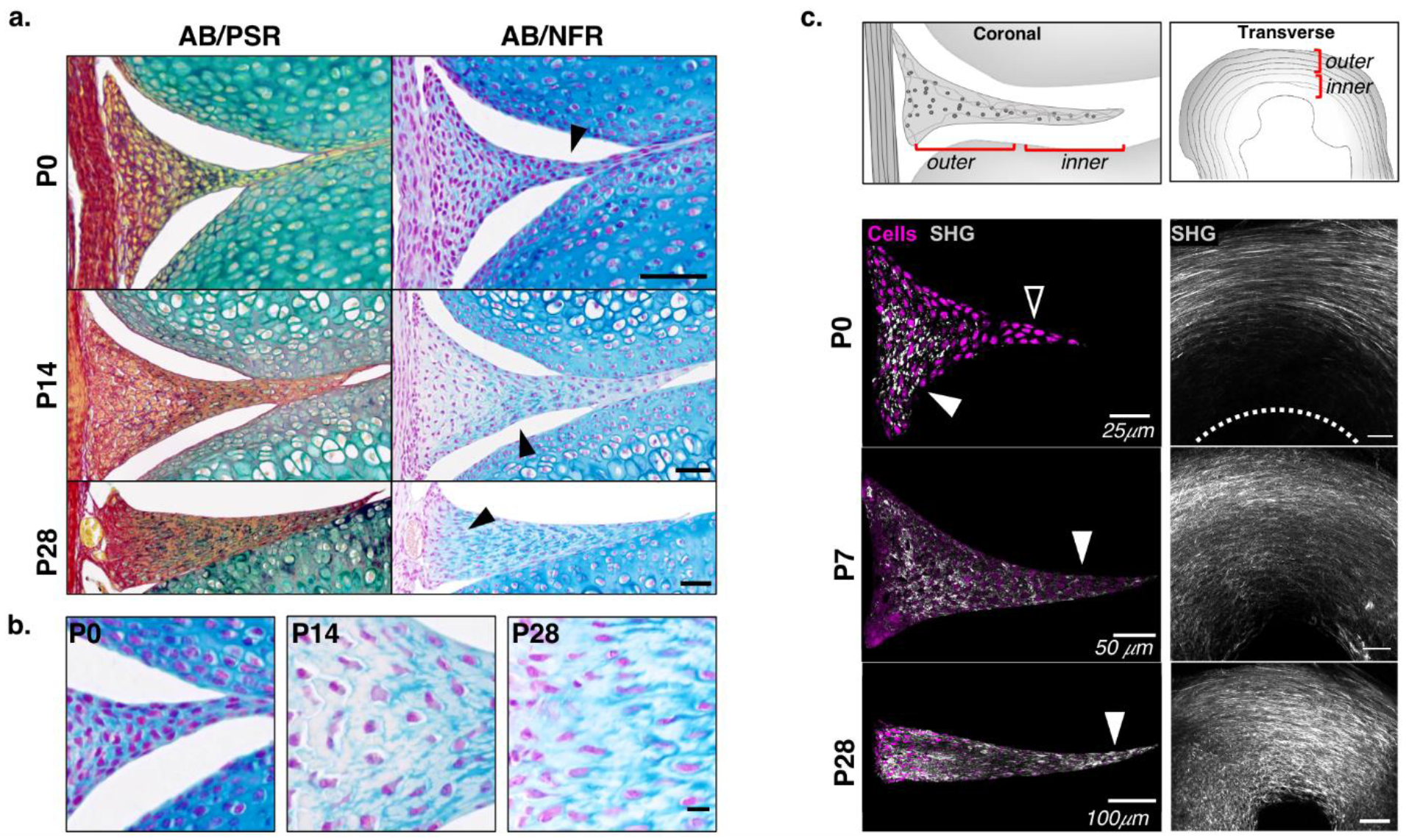
Distribution and organization of matrix components shift rapidly with age. **a)** Representative histological stain images of P0, P14, and P28 menisci showing co-distribution of collagen (Picrosirius Red, PSR) and proteoglycan (Alcian blue, AB) matrix components, and adjacent sections stained for AB and counterstained with nuclear fast red (AB/NFR) to show the localization of the proteoglycan matrix around cells. Scale bar: 50μm. Black arrowheads indicate location of the inset panels shown in **b)**. **c)** Second harmonic generation imaging (SHG) for visualization of fibrillar collagen matrix in coronal sections (left) and dissected menisci imaged in the plane of the circumferential fibers (right) at P0, P14, P28. White dashed line in the right P0 panel indicates the inner tissue boundary. An absence of SHG signal is noted within the inner region of the P0 meniscus (left: blank arrowhead, right: region proximal to white dashed line) compared to the outer region (left: white arrowhead, right: circumferential outer fibers). SHG signal within the inner region of the tissue appears at P7 and P28 (white arrowheads). Due to the extreme changes in aligned collagen content, laser settings were not kept equivalent between ages for visualization purposes, such that pixel intensity scales are not equal between the images.

### Distinct regional transcriptional profiles become more distinct during meniscus maturation

Our detailed analysis of tissue growth and ECM accumulation revealed both innate and maturation driven distinctions in the inner and outer meniscus cellular microenvironments. To explore how changes in resident cell phenotype beget these compositional distinctions, we compared the gene signatures of inner and outer meniscus cells to that of cartilaginous and ligamentous cells as a function of age. To do so, we used laser capture microdissection (LCM) to isolate inner and outer meniscus tissues (IM and OM), as well as adjacent articular cartilage (AC) and medial collateral ligament (MCL), from P0, P14, and P28 knee joint sections. Given the fibrocartilaginous nature of the meniscus, incorporating the AC and MCL into our analysis provided cartilaginous and ligamentous tissue benchmarks against which meniscus tissue regions could be compared. Using a custom microfluidic qPCR array, we assessed the expression of 93 select genes (representing collagens, proteoglycans/glycoproteins, ECM-remodeling factors, connective tissue transcription factors, cell-ECM interaction proteins, and mechanotransduction and nuclear envelope components, see **Supplemental Table 1**) (**Fig. 3a**).

**Figure 3:**
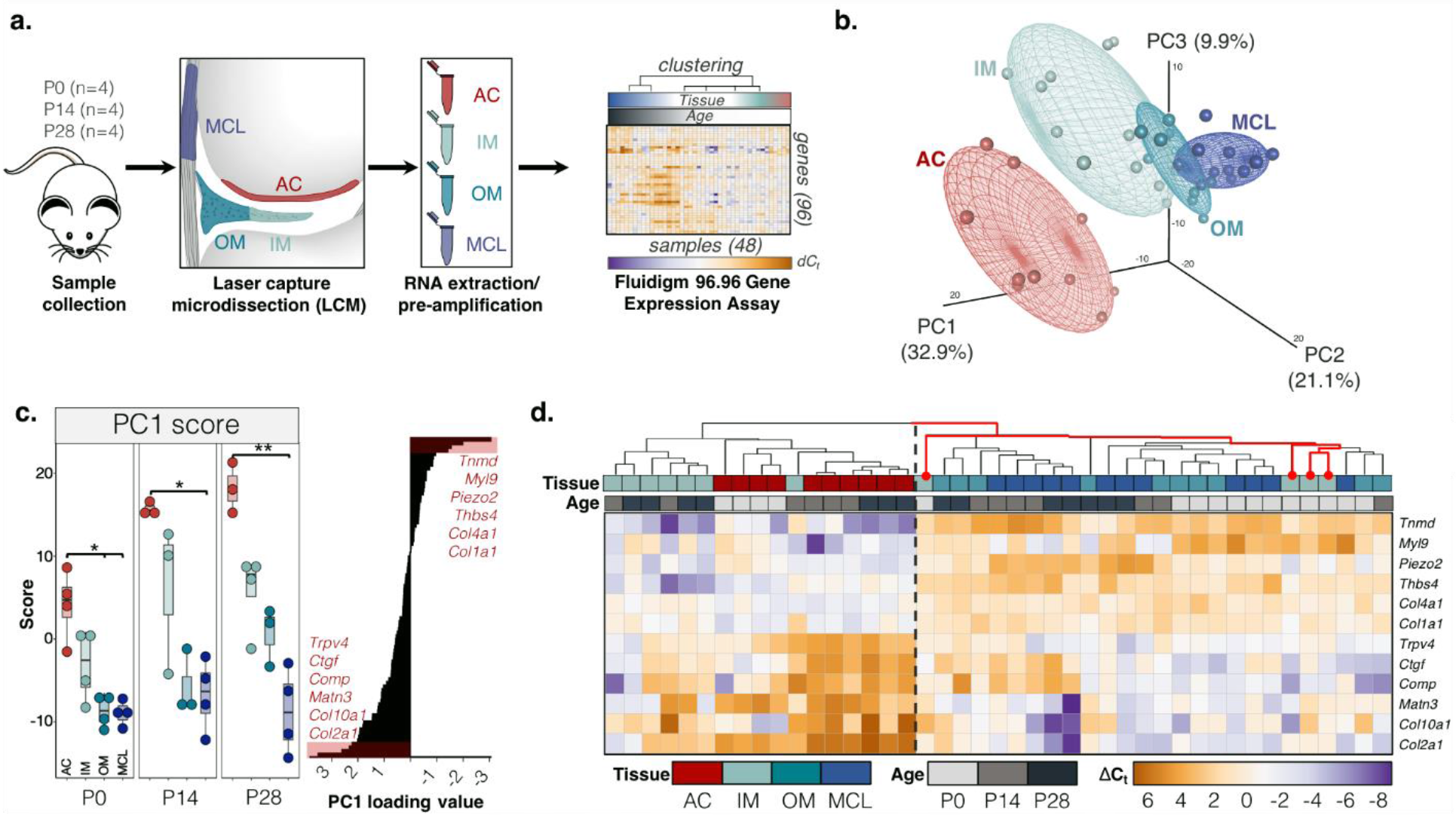
Comparison of fibrous and cartilaginous expression profiles across the knee joint during postnatal growth. **a)** Schematized approach to gene expression analysis (full procedure in methods). Articular cartilage (AC), outer meniscus (OM), inner meniscus (IM), and medial collateral ligament (MCL) for P0, P14, and P28 animals (n=4 per time point) were isolated for each knee joint of each animal via laser capture microdissection (LCM). Expression of the selected 96 genes (Supplemental Table 1) was measured using the Fluidigm 96.96 gene expression array, and is reported as delta-Ct (ΔC_t_) for each sample. **b)** Principal component analysis (PCA) of the data represented by mapping principal component scores for tissues of all age groups along the major principal component (PC) axes. Percent of described total variance by each PC is listed in parentheses. Concentration ellipses for each tissue are shown via clustering. **c)** (Left) PC1 scores compared for tissues at each time point. Kruskal Wallis analysis with a Dunn post-hoc test with Bonferroni p-value correction was used to statistically compare groups. *: p<0.05, **: p<0.01. (Right) Ranked PC1 loading values for analyzed genes, listing the 6 highest (*Col2a1* highest on list) and 6 lowest (*Tnmd* lowest on list) valued genes. **d)** Heatmap of ΔC_t_ values for genes listed in **c)** along with the resultant hierarchical clustering based on all measured genes (86 total, after sample exclusion due to technical errors), color coded by tissue type and age, as denoted at the top heatmap. Dashed vertical line indicates the split of samples based on first clustering event. For the heatmap with a full gene list used in the PCA/cluster analysis, see Fig. 3 Supplement 2a.

Principal component analysis (PCA) of the dataset revealed that the first three components accounted for ∼64% of the total variance across tissue and age, and mapping of these principal components (PCs) showed the largest delineation was between articular cartilage and the rest of the tissues (**Fig. 3b**). Genes with some of the lowest (*Tnmd*, *Thbs4*, *Col1a1*) and highest (*Col2a1, Col10a1, Trpv4*) PC1 loading values were genes typically associated with ligament and cartilage, respectively (**Fig. 3c, Fig. 3 S1a,b**) (Clark *et al*., 2010; Caceres, Pfeifer and Docheva, 2018). In general, established ECM markers (*Col1a1, Col2a1, Acan, Prg4, Tnmd)* followed expected patterns, with *Col1a1* and *Tnmd* being more highly expressed in the meniscus and MCL than in articular cartilage, while *Col2a1*, *Acan*, and *Prg4* showing the opposite trend (**Fig. 3 S1**), at all ages. These data confirmed that our analysis could detect the divergence of tissue phenotypes. Hierarchical clustering also confirmed that AC samples, at all timepoints, clustered away from the MCL. The meniscus, on the other hand, fell between these two extremes, depending on zone and age (**Fig. 3d, Fig. 3 S2a**). Specifically, the outer meniscus (OM) grouped with the MCL at all ages, while the inner meniscus (IM) initially clustered with the MCL and OM at P0 (**Fig. 3d**, red lines), but shifted towards the AC cluster as the tissue matured (P14, P28). When we ran the cluster analysis at each age (thus removing age as a variable), we saw that the IM shifted increasingly toward AC with age (**Fig. 3 S2b)**. This transition in the IM transcriptional profile supports the idea that the inner meniscus gains a more cartilage-like phenotype as the tissue matures and suggest that P14 is the age at which this transition occurs.

The initial PCA and hierarchical clustering indicated a shift in phenotype of inner meniscus cells with age, prompting us to further investigate this regional specialization. We conducted PCA on just the inner and outer meniscus samples to eliminate the possibility that the outcomes were skewed by the difference of the AC samples from the rest of the dataset. PC1 scores significantly differed between the IM and OM at all timepoints, indicating that PC1 captured the variation between the two regions with age (**Fig. 4a**). As before, genes associated with ligament and cartilage tissues appeared at the extrema of the meniscus-specific PC1 loading values, with *Col2a1, Col10a1, Ctgf* having the highest scores and *Myl9, Thbs4, and Tnmd* having the lowest scores. Importantly, only P0 IM samples clustered with the OM samples, while 2 of 3 P14 and all of the P28 inner meniscus samples grouped separately–further emphasizing the emergent differences between the inner and outer meniscus with age (**Fig. 4a,b**).

**Figure 4:**
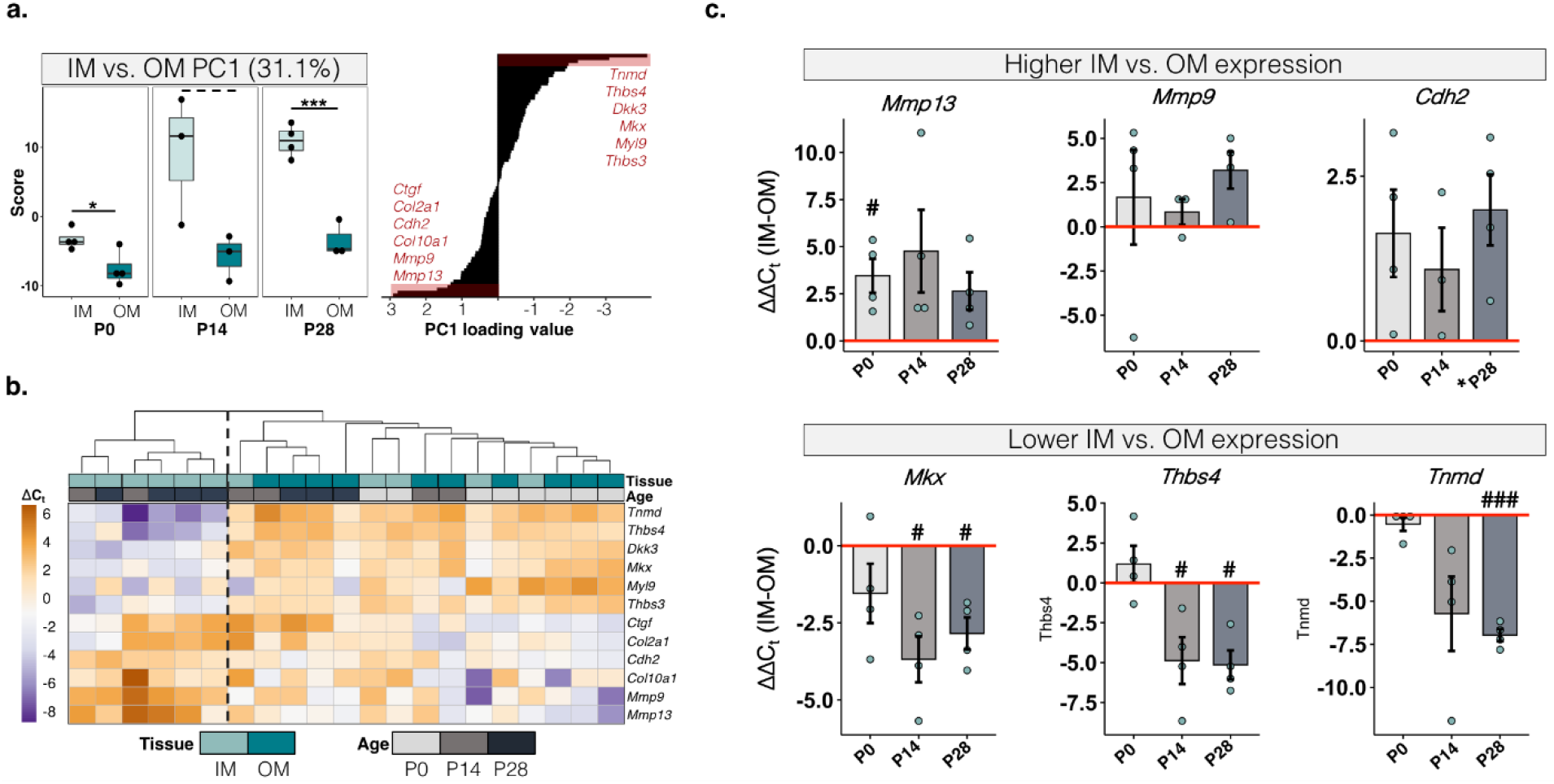
Gene expression signatures show specialization of the inner and outer meniscus during postnatal growth. **a)** PC1 scores for a PCA run on just the inner (IM) and outer (OM) meniscus samples. *: p<0.05; ***: p<0.001 by Student’s t-test. Dashed line indicates trend towards significance (p=0.09). **b)** Hierarchical clustering of IM and OM P0, P14, and P28 samples, as in Fig. 3d, with the separation of samples by the first clustering event denoted by the dotted line. Heat map represents ΔC_t_ values for the highest and lowest scoring PC1 genes from **a)**. **c)**ΔΔC_t_ values, ((ΔC_t_ of IM)-(ΔC_t_ of OM) for the same sample) for genes with higher IM expression relative to OM(top) and visa versa (bottom). Red line denotes no difference in expression (0). Mean ± sem shown. # indicates a significant deviation of the ΔΔC_t_ from 0 (i.e. a difference between the IM and OM ΔC_t_ values), as assessed by a one-sample t-test, #:p<0.05; ###:p<0.001.

Although an array of gene/molecule categories were included in the analysis (**Supplemental Table 1**), the largest differences in the inner to outer meniscus expression profiles were mostly in well-established ECM constituents (**Fig. 4c**). Interestingly, the inner meniscus had higher expression of several matrix metalloproteinases (*Mmp13*, *Mmp9*), with *Mmp13* being elevated in the inner relative to the outer region, even at birth, and both MMPs were enriched in this region by P28. N-Cadherin (*Cdh2*) expression was ∼4-fold elevated in the inner regions relative to the outer zones of these same samples. The inner region also had much lower levels of classic ligamentous markers that were abundant in the outer meniscus, including *Tnmd, Thbs4,* and *Mkx* (*Mohawk,* a transcription factor associated with ligament and tendon growth and development (Ito *et al*., 2010; Liu *et al*., 2010)). In many cases, these differences became more apparent with age, and once again pointed to a potential adaptation of resident meniscus cells to a differentially maturing extracellular matrix and loadbearing environment.

### Fluorescent collagen reporters highlight single-cell spatial heterogeneity in expression during regional specialization of the meniscus

While expression analysis revealed divergence of meniscus regions throughout maturation, this approach only surveyed the tissue at two week increments and used bulk measurements of all cells within selected regions. To increase the resolution of these spatiotemporal shifts in phenotype, we next employed a transgenic fluorescent reporter mouse model in which the expression of YFP, CFP, and mCherry fluorescent proteins are driven by the *Col1a1*, *Col2a1*, and *Col10a1* promoters, respectively (see methods). As the inner and outer regions of the mature meniscus have traditionally been described as Col II and Col I-rich, respectively, this model provided a novel way to examine the establishment of the expression profiles of these canonical ECM components at the single-cell level (Dyment *et al*., 2015). Broad assessment of fluorescence throughout the knee joint during maturation showed expected patterns, with *Col1*-YFP present in the ligaments and bone, *Col2-*CFP within the cartilage, and *Col10*-mCherry in the hypertrophic regions of tissues (**Movie 4**). Within the meniscus horns, there were clear nodules of Col-2 expressing cells transitioning to Col-10 expression, consistent with previous descriptions of ossification in this region (**Fig. 5 S1**)(Gamer, Xiang and Rosen, 2017). Notably, and unlike the horns, the meniscus body did not undergo ossification, even at adulthood. Instead, cells within the body of the meniscus exhibited a speckled, heterogeneous pattern of reporter expression (**Fig. 5 S1**, **Movie 4**). To determine whether this cell-to-cell heterogeneity differed with age and between inner and outer regions, we used the sectioning plane specified in **Fig. S1a** to define a coordinate system in which the position along the length of the meniscus is bounded by the inner (0) and outer (1) edge of the tissue. Cell position along the defined axis, as well as average *Col2*-CFP and *Col1*-YFP intensities in each cell, were then determined in samples taken from E17.5 to 4 months of age (**Fig. 5a**). This allowed us to (1) track how single cell *Col1a1* and *Col2a1* expression changed as a function of position within the meniscus (**Fig. 5b**), (2) evaluate co-expression of these two collagen reporters in individual cells (**Fig. 6a,b**), and (3) use these co-expression patterns to define distinct cell subpopulations across the meniscus including: negative (no reporter expression), *Col2*-CFP+ (only Col2-CFP expression), *Col1*-YFP+ (only Col1-YFP expression), and CFP+/YFP+ (co-expression of the reporters) (**Fig. 6c, Fig. 6 S1b,c**).

**Figure 5:**
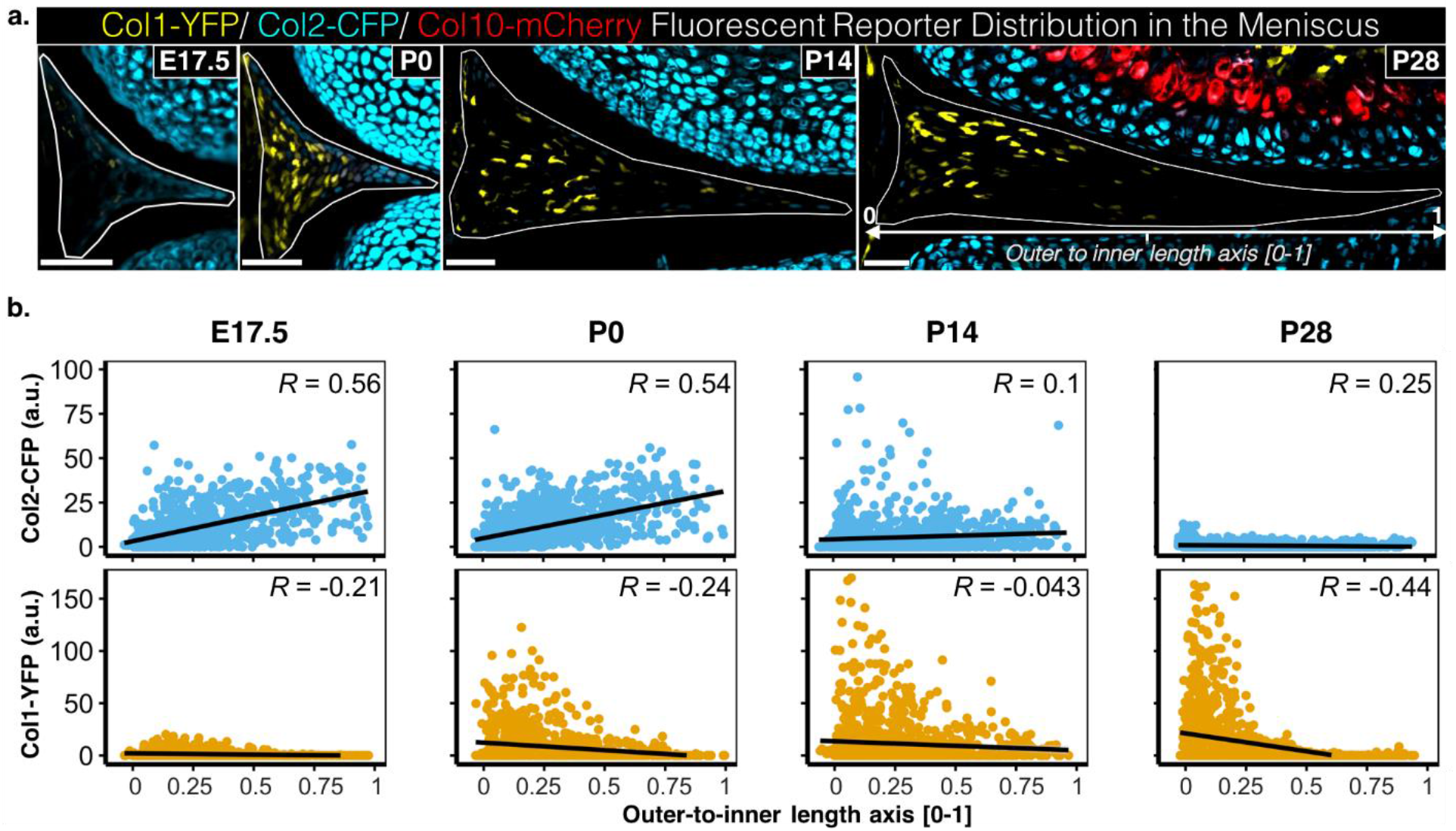
A Col-1/Col-2/Col-10 fluorescent reporter transgenic mouse allows for single-cell spatiotemporal tracking of the meniscus cell populations. **a)** Representative coronal sections from the meniscus body at E17.5, P0, P14, P28, showing the cellular expression of YFP, CFP, and mCherry fluorescent proteins driven by the *Col1a1*, *Col2a1*, and *Col10a1* promoters, respectively. Scale bar: 50*μ*m. Cell location (used here and Fig. 6) is defined by the ‘outer to inner length axis’, annotated in the P28 image, where the outer–most edge is the origin (0), the inner-most edge is 1 and a position in between is reported as the fraction of distance to the inner edge. For visualization, fluorescence intensities were adjusted separately for each age (i.e. LUTs were not kept constant between shown images). **b)** Fluorescence intensities for CFP (Col-2 reporter, top panel) and YFP (Col-1 reporter, bottom panel) for individual cells as a function of each cell’s location along the outer-to-inner length axis. Black lines, R, and p-values shown for Spearman correlations. n =777 (E17.5); n=888 (P0); n=1105 (P14); n=1122 (P28), with cells pooled from analyses of medial menisci from 3 animals at each age. For details on imaging and quantification (here and Fig. 6), see methods.

**Figure 6:**
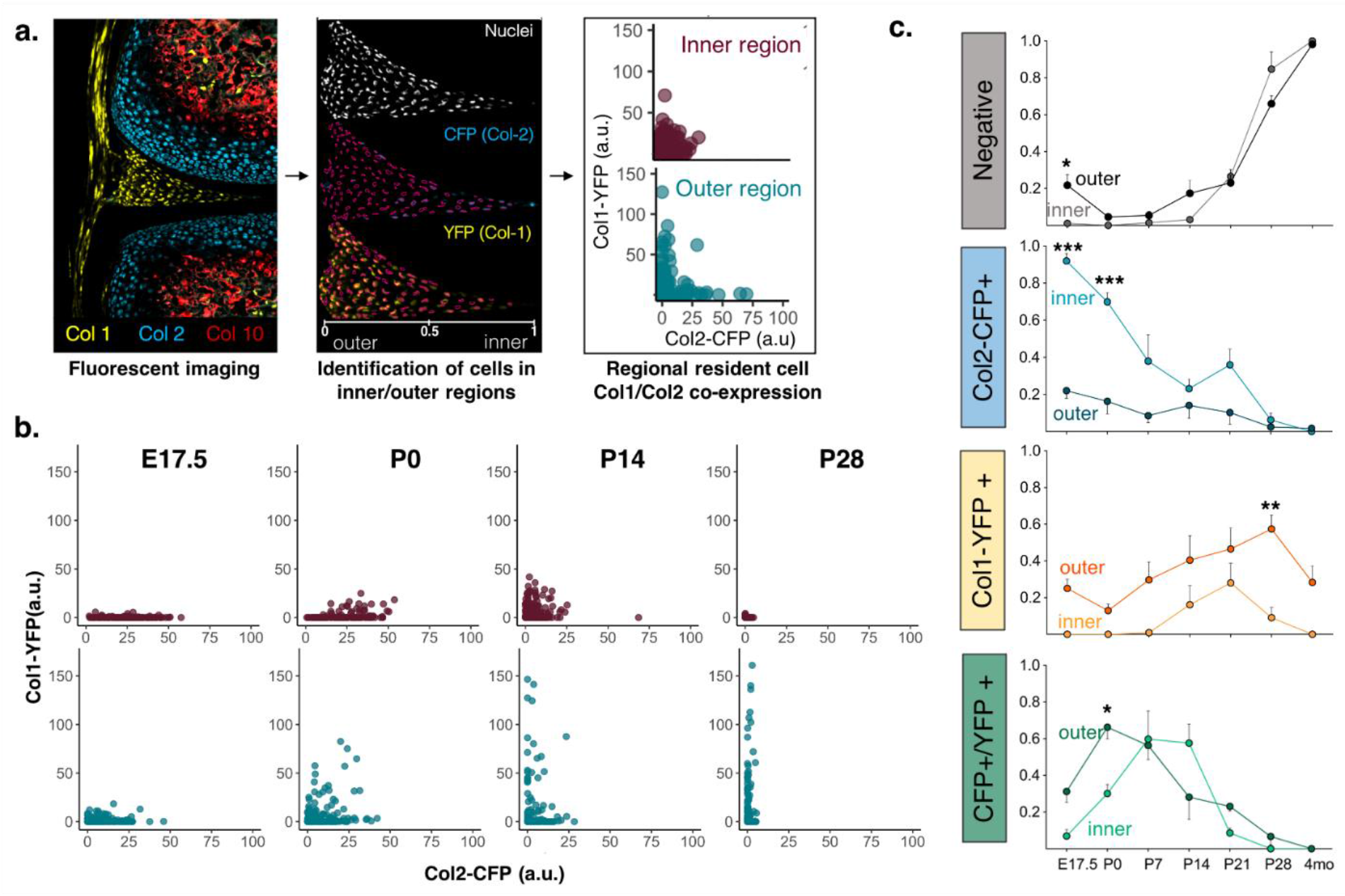
Col-1/Col-2 fluorescent reporters capture divergence of the inner and outer meniscus cell subpopulations in both early and late stage maturation phases. **a)** Workflow for single cell inner-to-outer quantification of *Col1*-YFP and *Col2*-CFP reporter co-expression through nuclear segmentation and quantification of fluorescence intensity for cells in the inner third (inner) and outer third (outer) regions (see methods for a detailed description). **b)** Fluorescence intensities of YFP (Col1 reporter) and CFP (Col2 reporter) for individual cells in the inner third (top row) and outer third (bottom row) of E17.5, P0, P14, and P28 menisci. **c)** Percent of cells in either the inner third or outer third of the meniscus that are classified into one of the following subpopulations based on their YFP and CFP fluorescence expression (see Figure 6 Supplement1 for details): 1. Negative (no detected reporter expression), 2. CFP+ (only the *Col2*-CFP reporter expressed), 3. YFP+ (only the *Col1*-YFP reporter expressed), or 4. CFP+/YFP+ (both reporters expressed). Shown as mean ± sem from 3 biological replicates. * (p<0.05), ** (p<0.01), ***(p<0.001) indicate significant difference between inner and outer regions at specified timepoint as determined by Bonferroni post-test of a 2-way ANOVA.

In the newly formed meniscus (E17.5 and P0), *Col2*-CFP fluorescence intensity positively correlated with outer-to-inner position, while *Col1*-YFP expression showed a negative correlation. This suggested that cells with higher *Col2* and *Col1* reporter expression were already segregated to the inner and outer regions of the meniscus, respectively (**Fig. 5a,b**). Interestingly, between E17.5 and P0, a distinct subgroup of outer meniscus cells emerged that had substantially higher *Col1*-YFP fluorescence than cells of the inner meniscus (**Fig. 6a,b**). The outer meniscus in this E17.5-P0 timeframe also showed a decrease in the number of negative, *Col2*-CFP+, and *Col1*-YFP+ cells and an increase in the number of double positive (CFP+/YFP+) cells. These data suggested that the presence of higher *Col1*-YFP expressors at P0 was a result of a subpopulation of outer meniscus cells that previously either expressed only *Col2*-CFP, or had previously expressed no reporters and subsequently increased their *Col1*-YFP levels (**Fig. 6c**). Meanwhile, and consistent with the correlation data, the inner meniscus contained significantly more *Col2*-CFP+ cells, further highlighting that distinctions between inner and outer meniscus cells are present by birth (**Fig. 6c**). *Col1*-YFP fluorescence values in the inner region peaked at P7 (**Fig. 6 S1a,b**) before diminishing at all subsequent ages. Overall, less than 18% of cells were negative for reporter fluorescence throughout the early growth stages (E17.5-P14), underscoring that this maturation window is characterized by high transcriptional and synthetic activity (**Fig. 6c**).

Later stages of maturation (P14-P28) demonstrated a shift in these trends. A negative correlation between outer-to-inner distance and *Col1*-YFP expression was maintained through P28, indicating that higher fluorescence in the outer meniscus persisted with age (**Fig 5b**). A gradual restriction of the *Col1*-YFP positive cells to the outer third of the meniscus occurred during this timeframe, and this remained so through 4 months of age (data not shown) (**Fig. 5b, Fig 6a,b, Fig. 6 S1a,b**). The *Col2*-CFP fluorescence intensity, however, showed a dramatic attenuation between P14 and P28. This was distinct from the bulk LCM/qPCR measurements, which showed *Col2a1* C_t_ values remaining consistently high in both the inner and outer meniscus at all timepoints (**Fig. 4**, **Fig. 5b**, **Fig. 6b**, **Fig. 6 S1a**). The decrease in fluorescence was further reflected by the increase in the “negative” cell subpopulation, which reached more than 60% in both regions, with a minimal presence of *Col2*-CFP+ and CFP+/YFP+ cells. As expected, however, high *Col2-*CFP fluorescence persisted in cells of the articular cartilage, suggesting a potential ‘threshold’ of *Col2a1* expression at each age above which CFP fluorescence can be detected (**Movie 4**, **Fig. 5a**, **Fig. 6 S1c**). Despite this limitation, the reporter expression analysis as a whole identified cell-to-cell heterogeneity present during the maturation process. These data further support that inner/outer regional cellular subpopulations are present as soon as the meniscus is formed, and further segregate and specialize at later maturation timepoints.

## DISCUSSION

The impressive strength and durability of mature fibrous tissues and their extremely limited tissue turnover and regenerative capacity underscores the importance of proper ECM assembly during postnatal growth. The knee meniscus offers a particularly interesting case of ECM design, wherein the existence of regionally distinct matrix organization and composition is appreciated as functionally important. Yet, the initial stages of specification and assembly that lead to the establishment of a proteoglycan-rich inner zone resisting compressive loading and an aligned collagenous outer zone resisting tensile loading within this tissue are poorly understood. Therefore, this study tracked the growth of the murine meniscus from its embryonic formation through its first month of growth—capturing a critical time-window during which animals begin to ambulate and weight bear. Interestingly, we found that there were distinct matrix and cellular features that defined the inner and outer meniscus at birth, indicating that the act of bearing weight during ambulation itself does not appear to be critical to cellular specification in these tissue regions. We also demonstrate, however, that zonal differences were further refined with postnatal growth; highlighting that the complex, multi-axial loading environment of the knee is likely critical in region-specific cellular specialization and functional matrix assembly.

Our analysis demonstrated that by birth, the classic circumferential fibrillar collagen alignment in the meniscus was evident, but was localized to just the outer region of the tissue wedge. Likewise, the inner region was already enriched in proteoglycans and devoid of detectable fibrillar collagen content (**Fig. 2, Movie 1**). These differing matrix characteristics suggest that synthetic activities of resident cell populations diverge during embryonic stages of meniscus development, prior to substantial extrinsic forces from weight bearing. This in turn creates distinct microenvironments that may further influence the phenotype of residing cells. In support of altered synthetic activities, fluorescent reporter analysis showed phenotypic divergence of inner and outer cells as early as E17.5—a few days before birth. Localization of cells with higher *Col2*-CFP reporter expression to the inner portion of meniscus and *Col1*-YFP increased specifically in outer meniscus cells between E17.5-P0 indicated a region-specific shift in cell behavior during embryonic growth (**Fig. 5; Fig. 6b; Fig. 6S1b**). Using LCM to specifically isolate inner and outer meniscus cells from the same tissue section, we were able to confirm discernable gene signatures in the inner and outer meniscus tissues at P0 (**Fig. 4b**). Expression of matrix proteases *Mmp13* and *Mmp9* was enriched in the inner meniscus, suggesting that the region may be devoid of fibrillar collagen due to heightened matrix remodeling activity. Conversely, decorin (*Dcn*)—a protein involved in collagen fibrillogenesis—and an aggrecan protease (*Adamts4*) were expressed at higher levels in the outer region, indicating that outer cells may be producing matrix components that promote fibrillar matrix assembly while resorbing proteoglycans in the region (**Fig. 4c**, **Fig. 4S1**). Of note, higher expression of MMPs and decorin in the inner and outer portions, respectively, persisted throughout maturation, indicating that differential expression of certain matrix proteins may be intrinsic to the inner and outer cell populations. In all, our data demonstrate regional meniscus cell distinctions as early as the embryonic stages of meniscus growth. This finding raises the question of whether these cells are derived from separate developmental origins or if the early establishment of mechanical and topographical differences may drive cell behavior in these regions based on physical inputs and timing of condensation within the meniscus structure (Hyde, Boot-Handford and Wallis, 2008; Shwartz *et al*., 2016). Further study of meniscus embryonic development, which includes rigorous lineage tracing and assessment of the biophysical properties of the nascent tissue, is required to confirm or refute these hypotheses.

In addition to identifying distinction in cell and matrix at birth, this work also sheds light on the substantial morphologic and cellular evolution that the meniscus undergoes during postnatal growth and maturation. Our tracking of the growth of the meniscus body confirmed previous findings (in both small and large animal models) showing that tissue growth is characterized by rapid accumulation of fibrous, collagen-rich matrix throughout the length of the tissue (**Fig. 1, 2**)(Gamer, Xiang and Rosen, 2017). We demonstrate that in mice, however, the majority of this postnatal growth occurs in the first two weeks of life and does not happen isometrically. Rather, the inner region of the tissue appears to extend inward to cover the widening tibial plateau (**Fig. 1 b-d**). It is interesting to consider whether this tissue lengthening during maturation is the result of directed matrix deposition (guided by the concurrent widening of the femoral and tibial surfaces), or if these postnatal shape changes represent an adaptation that increases efficiency of load transfer as animals begin to weight bear. Consistent with humans, the percent coverage of the meniscus on the tibial plateau (**Fig. 1d**) increases with age, suggesting that the latter mechanism may be likely (Clark and Ogden, 1983). Likewise, the appearance of proteoglycan-rich matrix in the pericellular spaces from P14 onwards highlights an alteration of matrix protein production in resident cells; possibly as an adaptation to increased compressive loading (**Fig. 2**).

Coincident with this matrix remodeling, gene expression patterns also evolved between the inner and outer meniscus throughout postnatal maturation (P0-P28). Importantly, our assessment of cellular phenotype (by qPCR and fluorescent reporters) was performed relative to the adjacent ligamentous (MCL) and cartilaginous (AC) tissues within the same knee joints at the same time (**Fig. 3**, **Fig. 6**, and supplements). Using these tissues as benchmarks of canonical ECM structures acting in tension and compression, respectively, further contextualized the maturation-driven divergence of cell populations in the meniscus regions. Specifically, inner meniscus cells converged to a more cartilage-like transcriptional profile by P28, while outer meniscus cells took on expression patterns that were more ligament-like, indicating that the specialization of these cell types with maturation may indeed be the result of exposure to distinct types of forces (**Fig. 3**). In the outer region, this was particularly exemplified by the dramatic increase of *Tnmd*, a glycoprotein highly expressed in tendons and ligaments (Docheva *et al*., 2005; Shukunami *et al*., 2006), relative to cells of the inner region by P28. Concurrently, there was a reduction of aggrecanase (*Adamts4*) transcripts, perhaps explaining the pericellular PG accumulation in the region by this timepoint. The restriction of the *Col1*-YFP reporter to cells of the outer third of the tissue by P28 further supported this specialization of outer meniscus cells (**Fig. 5, Fig. 6b**). Notably, *Col1*-YFP fluorescence intensities were heterogeneous, with a distribution of low and high expressors that was most variable during the timeframe of rapid postnatal growth between P0 and P14 (**Fig. 6 S1b**). The single cell ‘snapshot’ of reporter expression may indicate transcriptional ‘burstiness’ in individual cells which, when summed, would faithfully reproduce the bulk transcriptional profiles we measured via qPCR. This may suggest that deposition of ECM components that establish the mature tissue structure is accomplished by subpopulations of ‘high expressers’ of various matrix components rather than a uniform synthetic contribution from the resident cells. In fact, we’ve identified other fluorescent lines, with reporters for *Col6a1, Scx,* or *Dkk3*, that label distinct subsets of meniscus cells (**Fig. 5S1b**) and can further illuminate the spatial characteristics of the resident cell subpopulations. In combination with analyses like single cell RNA seq, these genetic tools may offer unique ways of not only establishing how cellular heterogeneity contributes to fibrocartilage assembly, but also studying its role in the injury and disease of these complex structures.

While our work demonstrates that assembly of the mature fibrocartilage tissue occurs through a synergistic system of increasingly specialized cell subpopulations and accretion of ECM, how much of the observed changes in meniscus cell phenotype is a direct response to the rapidly changing microenvironment (i.e. cellular interpretation of the extracellular biophysical cues) rather than differences in cell fate remains an important area of study. Indeed, extensive work in 2D and 3D *in vitro* systems has established mechanobiologic principles regarding how biophysical cues such as cell-cell/cell-ECM contacts, matrix topology, matrix stiffness, and applied external forces are sensed and interpreted by cells (Engler *et al*., 2006; Kumar, Placone and Engler, 2017). The presence of distinct matrix organization at P0, coupled with increasing extracellular fibrillar collagen content (**Fig. 2**), decreased cellularity (**Fig. 1**), and postnatal limb movement, all point to meniscus cells experiencing changes in key biophysical inputs throughout maturation, especially in the first two weeks of rapid growth (E17.5-P14, **Fig. 1**) (McNulty and Guilak, 2015; Qu *et al*., 2018). In fact, previous AFM measurements of murine meniscus ECM indicate a substantial stiffening of the matrix with age, another important parameter that may influence meniscus cell phenotype (Sanchez-Adams and Athanasiou, 2012; Sanchez-Adams, Wilusz and Guilak, 2013; Li *et al*., 2015, 2017). Interestingly, however, though we incorporated cell mechano-sensing factors as part of our gene expression analysis (contractility proteins, cell-cell/cell-ECM contracts, mechanosensitive transcription factors), it was the classical musculoskeletal markers (ECM proteins, connective tissue transcription factors) that distinguished inner and outer regions across growth (**Fig. 3c, Fig. 4a, Supp. Table 1**). This may suggest that the machinery required to interpret mechanical cues in meniscus cells is already in place by birth. Furthermore, the finding that inner and outer cell populations are distinct at birth, and may actually already differ at E17.5, is an important one—highlighting the possibility that the meniscus is truly shaped by cells of different fates (Hyde, Boot-Handford and Wallis, 2008). Strikingly, studies of immobilized embryonic chick limbs have shown that cell condensates that form the nascent meniscus eventually dissociate without the physical cues provided by muscle contraction, leading to loss of the meniscus tissue (Mikic *et al*., 2000). These studies highlight that mechanical stimuli are an important factor in meniscus morphogenesis as well, but the extent to which mechanics can specify distinct tissue zones and whether and when other biological stimuli are required, warrants further investigation.

Ultimately, uncovering the mechanism by which evolving biophysical cues (such as cellular organization, matrix composition and alignment, and extrinsic mechanical forces) influence the behavior of cells and the subsequent assembly of mature meniscal fibrocartilage requires a system that can be genetically and physically perturbed and can be studied throughout the critical timepoints of embryonic development and early maturation. The high spatiotemporal resolution of our analysis provides a strong foundation for future work focused on meniscus matrix assembly by creating a timeline of the concurrent establishment of both matrix properties and cell phenotype throughout maturation. Based on these critical findings, future work perturbing key mechanical inputs such as matrix organization, stiffness, and limb movement during time windows of murine maturation defined by this study will be able to shed light on mechanobiologic mechanisms associated with the assembly of composite ECM structures.

## MATERIALS AND METHODS

### Animals

All animal housing, care, and experiments were performed in accordance with the UPenn IACUC (protocol# 806669). All maturation experiments were done using the Col1-YFP/Col2-CFP/Col10-mCherry triple transgenic line (Maye *et al*., 2011; Dyment *et al*., 2015; Utreja *et al*., 2016). These CD1 background strain mice contain 3 transgene insertions: (1) a 3.6KB fragment of the *Col1a1* promoter driving the expression of the YFP gene (Tg(Col1a1*3.6-Topaz)2Rowe)(Kalajzic *et al*., 2002), (2) *Col2a1* promoter driving the expression of the CFP gene(Tsumaki *et al*., 1999; Chokalingam *et al*., 2009; Maye *et al*., 2011), and (3) the *Col10a1* promoter driving the expression of the mCherry fluorescent protein gene (Tg(Col10a1-mCherry)3Pmay/J)(Maye *et al*., 2011). To provide examples of other fluorescent reporter mouse models that could be useful for meniscus research (**Fig. 5 S1b**), the Scleraxis-GFP (Pryce *et al*., 2007; Blitz *et al*., 2009), Col6a1-GFP (STOCK Tg(Col6a1-EGFP)JB13Gsat/Mmucd from MMRRC), and Col1-CFP/Dkk3-GFP (Utreja *et al*., 2016) transgenic mouse models were used.

### EdU labelling

For analysis of cell proliferation (**Fig. 1f**), animals were weighed and injected with 3 µg/g 5-ethynyl-2’-deoxyuridine (EdU) for two consecutive days prior to the P4, P14, and P28 sacrifice time points. Three animals per time point (1 limb/animal) were sectioned (see “Tissue sectioning”), stained using the CalFluor 647 Azide Kit (Click Chemistry Tools, Cat#: 1372), and imaged (see “Fluorescent tissue imaging/fluorescence image analysis”).

### Tissue harvesting

For tissue growth measurements, EdU labeling analysis, SHG imaging, and fluorescent reporter imaging, animals were euthanized, hindlimbs were disarticulated, and the skin was removed. Knee joints were fixed in 10% neutral buffered formalin for 2 days at 4°C, followed by a 1day incubation in 30% sucrose at 4°C, after which samples were embedded and frozen in OCT. N=3 animals/age (E17.5-4mo) was harvested for analysis. For laser capture microdissection/gene expression analysis, hindlimbs were harvested at P0, P14, P28, and quickly transferred to a 4% phosphate-buffered paraformaldehyde (PFA) solution and fixed at 4°C on a shaker for 2 hours. Samples were then quickly rinsed in ice-cold RNAse free PBS and embedded in OCT to minimize RNA degradation (n=4 animals/timepoint). For histologic analysis, tissues were fixed in 4% PFA for 2 days at 4°C. Samples were then decalcified in a 10% EDTA/2% PFA solution that was changed every other day. P0-P7 samples were decalcified for 4 days and P14-P28 samples for 14 days, prior to paraffin embedding using a tissue processor (n=3 animals/timepoint) <u>Tissue sectioning:</u> Knee joints were serially sectioned from the patellar tendon through the posterior medial meniscus horn in the coronal plane (**Fig. 1a**). Care was taken to maintain the sectioning plane consistent between samples to ensure accurate comparison between samples. Sections were also monitored to ensure that the medial and lateral anterior meniscus horns were in the same sectioning plane (proper medial/lateral alignment) and that the full length of the MCL was visible in more posterior regions of the knee (no anterior tilting of the femur/tibia). Sections containing the body of the medial meniscus were then selected from the serial sections based on the triangular shape of the tissue, and only samples where proper alignment could be achieved were used for further analysis (see **Fig. 5 S1a** for representative shape change). OCT-embedded Col1-2-10 fluorescent reporter (and EdU labelled) samples were cryosectioned using cryofilm 2C (Section-lab Co), at a thickness of 8µm (Dyment *et al*., 2016). Samples used for LCM analysis were collected (8µm) using the CryoJane tape-transfer system (Electron Microscopy Science, Cat#: 62800). Paraffin section thickness was 5µm.

### Histology

For each stain (**Fig. 2, Fig. 2 S1**), paraffin sections for each time point were stained together for consistency. For proteoglycan content assessment, Alcian Blue (1%, pH 2.5, Rowley Biochemical) Toluidine Blue (0.025% solution, Sigma 364-M) or Safranin-O solution (0.2%, Sigma S2255) were applied using standard protocols. Alcian blue stained sections were counterstained with either Nuclear Fast Red (Electron Microscopy Sciences, 26078-05), or Picrosirius Red (0.1%, Rowley Biochemical SO-674). For Safranin-O, a 0.05% Fast Green Solution (Fisher Scientific, F99-10) was used as a counterstain for collagen/fibrous tissue content. Slides were imaged using a Zeiss Axio Scan.Z1 slide scanner using a brightfield setting and a 20X objective. Only medial menisci were used for analysis.

### Fluorescent tissue imaging

Sections were fixed to glass slides using a Chitosan adhesive and the full knee joint was imaged using the Zeiss Axio Scan.Z1 slide scanner using a 10X objective and the Colibri 7 LED illumination source. For EdU imaging, 2-3 non-adjacent stained sections from the meniscus body were counterstained with Hoechst 33342 and imaged using the darkfield, DAPI, and Cy5 emissions filter channels. For Col1-2-10 fluorescent reporter sections, all sections were counterstained with TOPRO-3 (ThermoFisher, Cat#: T3605). Samples from each age (E17.5-4mo) were imaged at the same time, with the same image settings (LED intensity, exposure times) to ensure fluorescent intensity comparisons between ages could be made. Darkfield, CFP, YFP, RFP, and Cy5 channels were imaged (to define tissue boundary, Col2 reporter, Col1 reporter, Col10 reporters, and nuclei, respectively). For all samples, at least 3 sections/knee joint were imaged for analysis, with no two sections adjacent to one another in the series of serial sections, to avoid analysis of the same cells (n=3 animals/age).

### Meniscus shape measurements

All image processing was performed with Fiji software (Schindelin *et al*., 2012). Darkfield and Cy5 (nuclei) channels of medial meniscus images were stacked, cropped, and rotated such that the edge of the tissue closest to the tibia was oriented parallel to the x-axis. Darkfield images were used to outline the meniscus boundary and the “Measure” tool used to determine the cross-sectional area. Meniscus length (M_L_) was determined by drawing a segment with the line tool parallel to the tibial meniscus edge from the inner to outer tip. Medial height (M_H_) was determined by drawing a line segment perpendicular to the M_L_ line segment from the outer point of the tibial edge of the tissue to the highest point in the meniscus outline (refer to schematics in **Fig.1**, and **Fig.1 S1a**). For tibial length, a line segment along the medial tibial plateau was drawn up to the site of the intra-articular ligament insertion, and length measured. To calculate % tibial coverage of the meniscus (**Fig.1d**), two perpendicular line segments to the tibial length segment were drawn: one that intersected the inner tip of the meniscus, and one that intersected the outer edge of the tibia. The distance between these two perpendicular lines was measured, and divided by the total measured length of the tibial plateau (**Fig.1 S1a**). Nuclei within the selected meniscus boundary were masked, and counted. Cellular density, within a section, was calculated as the number of counted nuclei per measured cross-sectional area. For inner and outer area and density measurements, the midpoint of the meniscus length line segment was used to split the cross section in half and outline the inner and outer area (**Fig. 1 S1a**). Area and cell density measurements were then repeated as described in the two regions.

For all measurements, 3 sections were measured and averaged per animal, and the average values for 3 animals per time point are reported.

### Second harmonic generation (SHG) imaging

All imaging was done using the Leica SP8 2-Photon Microscope with a 20X water immersion objective through the Penn Vet Imaging Core. For whole tissue imaging, the medial meniscus was dissected from formalin-fixed knees at indicated time points (**Fig. 2**), incubated in TOPRO-3 (1:1000 dilution in 0.02% Triton-X/PBS solution) for 2 hours on a shaker, and pressed between two No. 1 glass coverslips such that the circumferential fibers lay parallel to the imaging plain for optimal signal detection. A 2-photon laser (adjusted wavelength: 860nm) with activated forward scatter second harmonic generation (SHG) was used to simultaneously capture collagen fibers and stained nuclei throughout a 40-90µm tissue depth, with a step-size of 0.5-1µm. Laser intensity and gain was adjusted between ages to better the quality of the SHG signal for fiber visualization. For comparison of SHG signals between menisci of different ages, laser intensity was kept constant, and the tissues being compared were imaged on the same day (**Fig. 2 S1d)**. For SHG imaging of tissue sections, z-stacks (1µm z-height) of the 8µm sections prepared for fluorescence imaging (see below) were scanned using forward scatter SHG along with TO-PRO3 imaging of cells, with all imaging parameters (laser wavelength, power, gain) kept constant during imaging. Maximum intensity projections of the stacks are presented in **Fig. 2c, Fig. 2 S1c**. When appropriate, brightness and contrast settings were further adjusted in Fiji software for better signal visualization.

### Laser capture microdissection (LCM)

Sections (on CryoJane slides) were dehydrated through a ethanol gradient and dried immediately with xylene prior to laser microdissection. The Arcturus XT Laser Capture Microdissection (LCM) System (ThermoFisher Scientific) mounted on a Nikon Eclipse T*i* inverted microscope (10X, brightfield) was used to outline regions of interest for collection, microdissect the selected region using the Arcturus IR laser, and collect dissected tissues off of the slide using the CapSure Macro LCM caps (ThermoFisher, Catalog # LCM0211). For the meniscus, the midpoint of the coronal length (**Fig. 1 S1a**) was approximated, and served as the boundary between the inner and outer meniscus regions selected for collection (**Fig. 3a** schematic). For the articular cartilage (AC), only the superficial 3-4 cell layers (cells that remain non-hypertrophic throughout maturation, See **Fig. 5a**) were selected for collection at each age (P0, P14, and P28) and only from the medial femoral condyle adjacent to the medial meniscus. MCL samples were collected from the segment of the ligament connecting to the femoral head. See **Fig. 6 S1c** for schematic of tissue regions of the AC and MCL collected. After tissue collection, caps were checked under the 10X brightfield microscope, and any debris picked up during collection was removed by placing the cap on a light adhesive surface. Prior to collection, the average number of cells per region of tissue collected per section at each age was calculated, and the number of sections per animal per age for each tissue was varied such that the total number of cells collected for each tissue at each age for each sample was approximately equal.

### LCM tissue processing for gene expression analysis

*Digestion:* Film containing LCM microdissected tissues was detached from caps, and submerged in digest solution containing 1X Digest Buffer (Zymo Research, Cat#: D3050-1-5) and supplemented with 1mg/mL Proteinase K (Zymo Research, Cat#: D3001-2-5) for 1 hour at 55°C. Samples were then transferred to 65°C for 15 min. to further de-crosslink the sample.

*cDNA preparation and preamplification:* RNA isolation of digested samples was performed using the Quick-RNA MicroPrep kit (Zymo Research, Cat #: R1050). In-column DNAse-I treatment was performed for trace DNA removal before eluting RNA. Prior to cDNA synthesis, a second DNAse digestion was performed on the total eluted RNA volume (16µL) to remove gDNA using the ezDNAse kit (ThermoFisher, Cat#: 11766051) according to the manufacturer’s protocol. For cDNA synthesis, SuperScript IV VILO Master Mix (ThermoFisher, Cat#: 11756050) was used with 16µL of DNAse treated RNA elute for a 20µL reaction volume, and RT-PCR was run according to the manufacturer’s protocol. Generated cDNA was then preamplified for the specific 96 gene targets. A primer pool containing the 20X Taqman primers (FAM-MGB probes) of all assessed genes (see **Supplemental Table 1** for gene names and assay ID numbers) diluted 1:100 (0.2X) in DNA suspension buffer (10 mM Tris, pH 8.0, 0.1 mM EDTA) was generated. The Fluidigm Preamp MasterMix (PN#: 100-5580), primer pool, and DNAse free water were combined with 2µL of cDNA from each sample for a 5uL reaction volume. Samples underwent 15 cycles of preamplification according to the manufacturer’s protocol, and the resultant cDNA was diluted 1:5 in the DNA suspension buffer.

### Fluidigm gene expression chip and data post-processing

*Gene expression:* qPCR of LCM-collected samples was performed on the Fluidigm Biomark HD at the Penn Genomic Analysis Core using a Fluidigm Dynamic Array IFC (PN#: BMK-M-96.96) loaded with pre-amplified cDNA (see above) and 20X Taqman probes (full list of surveyed genes is indicated in **Supplemental Table 1**) for 48 samples and 96 genes. Brightfield images of the Fluidigm chip were used to detect bubbles in the reaction wells and excluded from the analysis. Sample measurements for *18S* (one of the 3 housekeeping genes) was excluded due to their low raw C_T_ values, indicating that cDNA levels were outside the dynamic range of the probe. *Habp2* and *Cdh3* were also excluded due to undetected measurements in ∼90% of the samples. In all, of the total 4608 reactions, 481 were excluded due to technical issues. To calculate ΔC_T_ values for genes for a given sample (**Fig. 3**, **Fig. 3 S1**, **Fig. 4 S1**), the C_T_ values of two housekeeping genes (Abl1, Rps17) were averaged, and subtracted for the measured C_T_ value of all the genes for that one sample. For ΔΔC_T_ calculations (**Fig. 4c**), the gene ΔC_T_ value for the outer meniscus for each sample was subtracted from ΔC_T_ value for that gene measured in the inner meniscus.

*Principal component analysis (PCA):* For PCA and hierarchical clustering, ΔC_T_ values for all genes (excluding the housekeeping genes) were put into ClustVis (Metsalu and Vilo, 2015). All pre-processing parameters, as well as the PCA of the dataset are saved in ClustVis under the settings ID: ZVWcdeVCNTbyVyJ. A threshold of 25% for missing values was set for both the rows and columns, which excluded *Acta2* and *Adamts5* from the gene set and excluded 5 samples from the 48 sample set. For PCA of just the inner and outer meniscus, AC and MCL samples were filtered out and similar thresholding applied in ClustVis (settings ID: KRukrmwSSrKohHE). All plotting was done using custom R scripts using ‘ggplot2’, ‘rgl’, and ‘Rcmdr’ packages.

### Fluorescent image analysis

Fiji software was used for all image processing and fluorescence quantification. The darkfield channel and selection tool was used to segment the medial meniscus boundary and remove any other tissue from the field of view. The nuclei were then identified via intensity thresholding and masking for each image. The ‘analyze particles’ function was then used to count cells in each section, and determine the mean intensity within the nuclear mask of each cell for the other fluorescence channels of interest (Cy5-Calfluor EdU or CFP, YFP fluorescent reporter).

For quantification of EdU labelling (**Fig. 1f**), a positive EdU signal fluorescence threshold was determined for each biological sample, the number of positive cells (in all sections for each sample) counted, and percentage (of total counted cells for all sections of the same sample) determined. The ratio corresponded to “% positive” EdU cells counted, and reported for each animal (n=3) at each time point (P4, P14, P28).

For characterization of *Col1*-YFP and *Col2*-CFP single cell reporter expression patterns (**Fig. 5, Fig. 6**), segmented meniscus images were rotated such that the tibial edge of the meniscus wedge was parallel to the x axis. The x coordinates of all nuclei were recorded and scaled to the total coronal length of the tissue, such that the inner-to-outer position of each cell was mapped (**Fig. 6a**). The total number of cells analyzed per time point was as follows: n=777 (E17.5), n=888 (P0), n=1234 (P7), n=1105 (P14), n=2148 (P21), n=1122 (P28), and n=888 (4 mo.). For analysis in **Fig. 5b** and **Fig. 6 S1a**, all the cells for all sections of all biological replicates were pooled and plotted. For inner and outer region analysis, cells with x coordinates between 0-0.33 (inner third) were analyzed as ‘inner’ and those with coordinates between 0.66-1 as ‘outer,’ to further segregate the two subpopulations. To assess variance of inner and outer cell CFP and YFP co-expression (**Fig. 6b**), n=140 cells for each region at each age were randomly selected for the pooled data set, to account for the difference in cell numbers between the inner and outer meniscus. This process was repeated 10 times to ensure repeatability of results (data not shown), and a representative subset is shown in **Fig. 6b**.

To categorize each meniscus cell as ‘negative’,’CFP+’,’YFP+’, or ‘CFP+;YFP+’ (**Fig.6c**, **Fig.6 S1d**), YFP and CFP intensities in articular cartilage (a control CFP+ and YFP-tissue) and MCL (a control YFP+, CFP-tissue) cells were measured for each section of each knee joint sample analyzed, using the same method as what is described for the meniscus. Meniscus cells within the section were then defined as follows: A CFP+ cell is one where the average CFP fluorescence intensity that is higher than the 3rd IQR of what was measured in MCL cells, and a YFP+ cell is one where the average YFP fluorescence intensity is higher than the 3rd IQR of what was measured in AC cells of that same section. If a cell was both CFP+ and YFP+, it was grouped into the double positive category. Conversely, if neither threshold was achieved, a cell was labeled as ‘negative.’ All analysis and plotting was done using custom R scripts using the ‘ggplot2’ package.

### Statistics

All growth parameters (n=3 biological replicates per age group) assessed in **Fig. 1** were assumed to be normally distributed and compared via 1-way ANOVA with Bonferroni pos-hoc analysis (alpha=0.05). For comparison of percent abundance of reporter subpopulations in the inner and outer meniscus with age (**Fig. 6c**), a two-way ANOVA with Bonferroni post hoc comparison was used to determine significant differences between inner and outer regions (alpha=0.5). All ANOVAs were performed in Prism 5 GraphPad commercial software.

For comparison of principal component scores and gene expression, the Shapiro-Wilkes normality test was used to determine the need for non-parametric testing. As such, all data (**Fig.3c, Fig. 3 S1, Fig.4a**) was analyzed using Kruskal-Wallis analysis of variance, with a subsequent Dunn’s Test for multiple comparisons (alpha=0.05) with a Bonferroni adjustment used to compare distinct groups. For **Fig. 4c**, gene ΔΔC_T_ differences IM samples compared to OM at each age were compared by one-sample t-tests (*μ* = 0, *α* = 0.05). All analysis was done using custom R scripts employing the ‘rstatix’ package.

## AUTHOR CONTRIBUTIONS

All authors contributed to the conception and design of the studies. TKT and XJ performed the experiments and analysis. All authors contributed to data interpretation. TKT, RLM, and NAD wrote the manuscript, and all authors edited the manuscript and approved the final submission.

## ACKNOWLEDGEMENTS

This work was supported by the National Institutes of Health (R01 AR075418, R00 AR067283, and P30 AR069619) and the Department of Veterans Affairs (IK6 RX003416). Additional support was provided by the Center for Engineering Mechanobiology (National Science Foundation, CMMI-1548571). We thank the UPenn Vet School Imaging Core (NIH grant S10 OD021633-01) for the use of the multiphoton microscope, the UPenn Center for Sleep and Circadian Neurobiology for the use of the ArcturusXT laser capture microscope, and the UPenn Genomic Analysis Core for running the Fluidigm GE 96.96 Dynamic Array.

## SUPPLEMENTAL FIGURES

**Figure 1 supplement 1:**
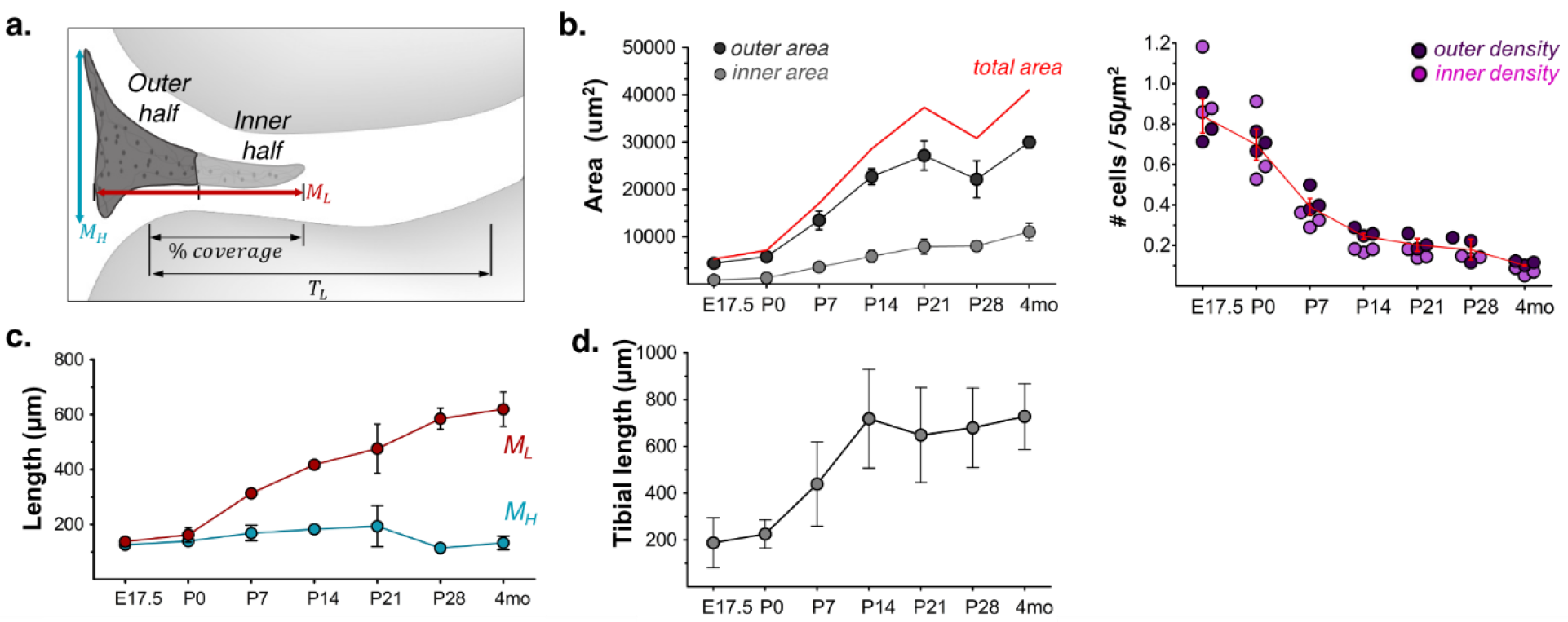
**a)** A more detailed annotation of meniscus growth measures, including segmentation of inner and outer meniscus area, and calculation of tibial length (T_L_) and percent tibial coverage by the meniscus. **b)**Areas of the inner and outer halves of the meniscus (as defined in panel a) with age. Total area (red line) from Fig.1 plotted for reference. Mean ± sd shown. **c)** Coronal length and height of meniscus sections with age showing a lengthening of the tissue. **d)** Total length of the medial tibial plateau with age. Mean ± sd shown.

**Figure 2 Supplement 1:**
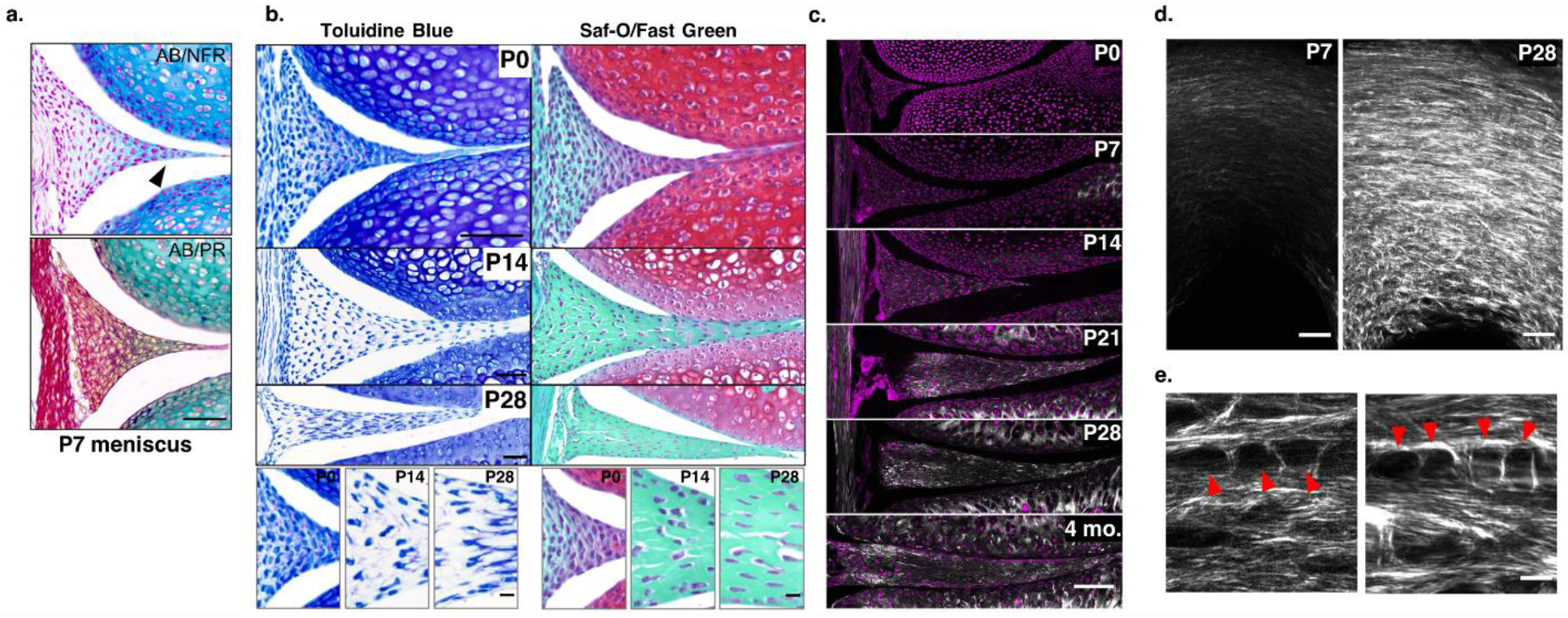
**a)** P7 serial sections stained for proteoglycans and nuclei (AB/NFR) or proteoglycans and fibrous matrix components (AB/PSR). Black arrowhead points to accumulation of proteoglycans in the inner region. Scale bar: 50μm. b) Serial sections of those shown in Fig. 2a, stained with Toluidine Blue (PGs in purple, nuclei in blue) or Safranin-O/Fast Green (PGs in red, fibrous matrix in green) as alternative indicators of proteoglycan-rich matrix. Inner region proteoglycan accumulation at P0 was detected by both stains. Pericellular accumulation at P14 and P28 could be confirmed via Toluidine Blue, but was not seen by Saf-O, indicating a difference in dye sensitivities. Scale bar: 50μm. **c)** SHG imaging (grey) with nuclear counterstain (magenta) of coronal meniscus body sections, as in Fig. 2b. Laser intensities/imaging acquisition settings were kept constant between ages to show progressive aligned matrix accumulation through an increase in SHG signal. Scale bar: 100μm. **d)** Maximum intensity projections through 30μm depth of a P7 (top, white outline denotes tissue boundary) and P28 (bottom) explanted meniscus imaged with the same acquisition settings (laser, gain, etc) to show the difference in SHG signal between the two samples. Scale bar: 50μm. **e)** Single plane SHG images from Movies 2 (P7) and 3 (P21) showing examples of fibrous lacunar structures in which cells reside (red arrowheads, cells not shown). Imaging settings were modified between the two ages due to large difference in SHG signal intensity.

**Figure 3 supplement 1.**
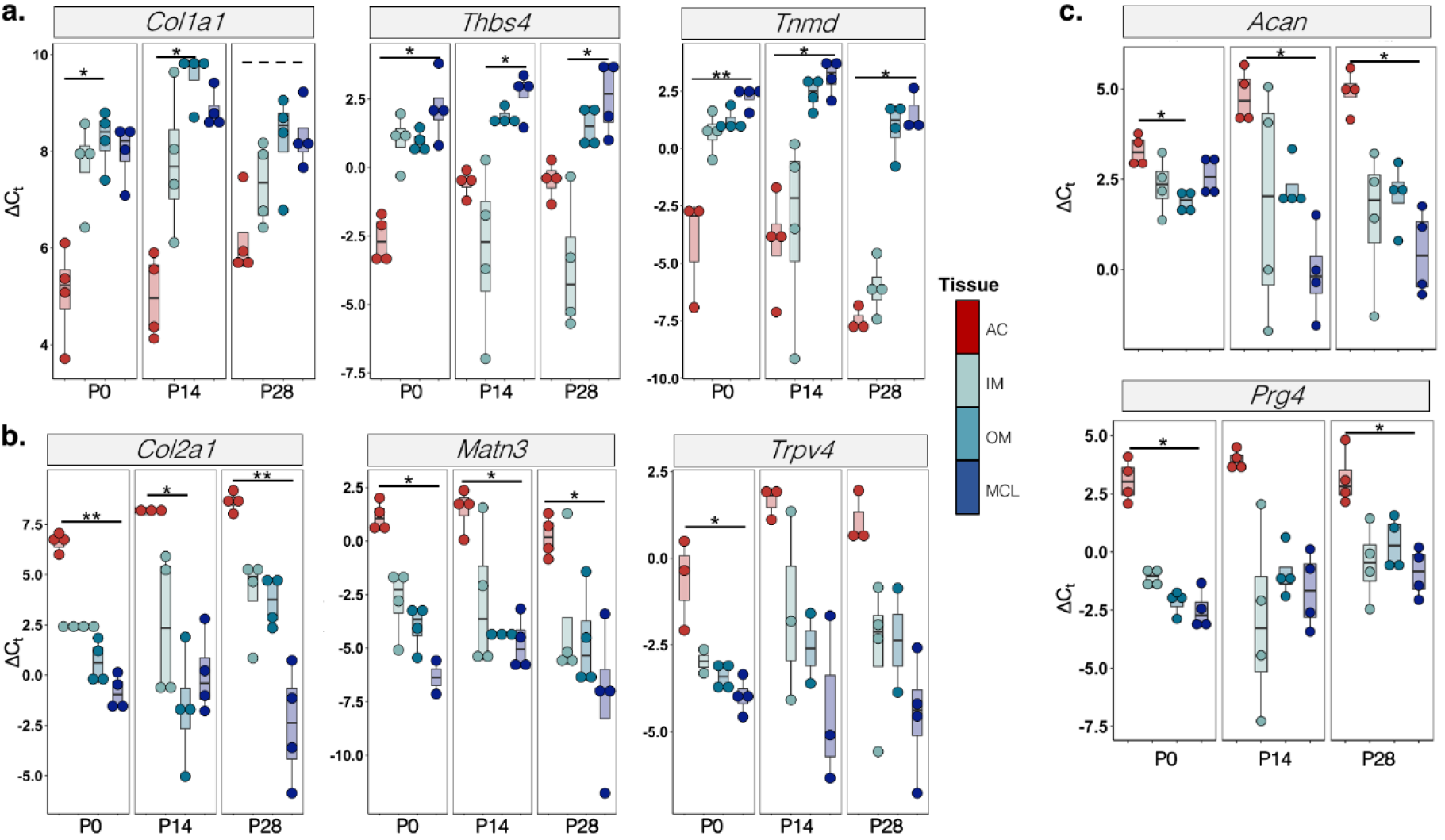
ΔC_t_ values of selected genes with the lowest (**a**) and highest (**b**) PC1 scores (as shown in Fig. 3c) for articular cartilage (AC), inner meniscus (IM), outer meniscus (OM), and MCL at P0, P14, and P28. **c)** Same gene expression data for an additional pair of canonical cartilage genes: *Prg4 (Lubricin),* and *Acan (Aggrecan*). Statistical differences determined by Kruskal-Wallis Test with Dunn’s multiple comparison test: *: p<0.05; **: p<0.01. Dashed lines indicate trends towards significance.

**Figure 3 Supplement 2:**
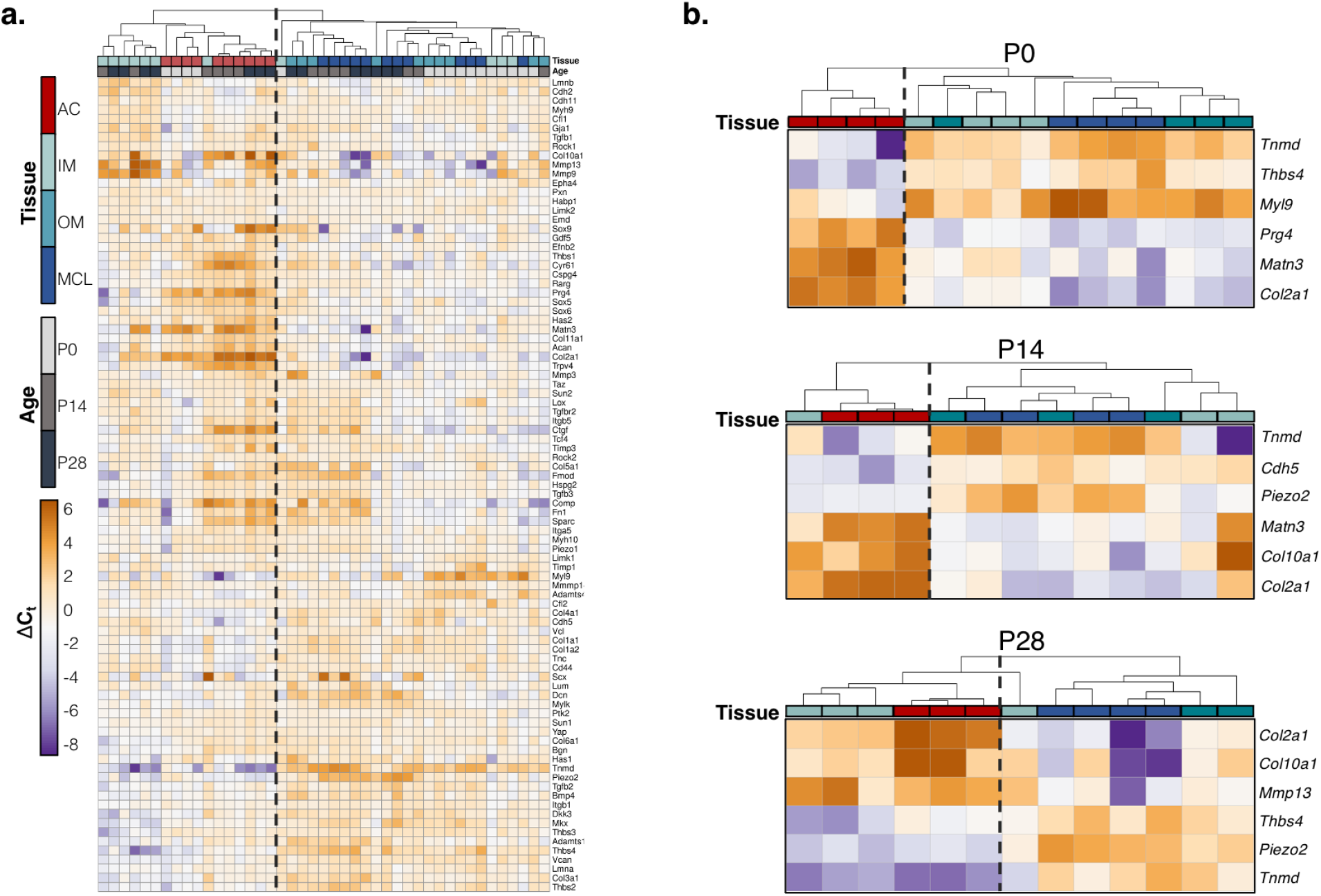
**a)** The full gene expression heatmap, showing clustering analysis as in Fig.3, as well as the entire analyzed gene panel. **b)** ClustVis cluster analysis performed at individual timepoints (P0, P14, P28), highlighting age-specific differences in tissue gene signatures. Genes on the heatmap represent the 3 highest and lowest PC1 loading scores based on PCA performed at each timepoint. The dashed vertical lines in **a,b** indicate the split of samples based on first clustering event (as in Fig. 3d).

**Figure 4 Supplement 1:**
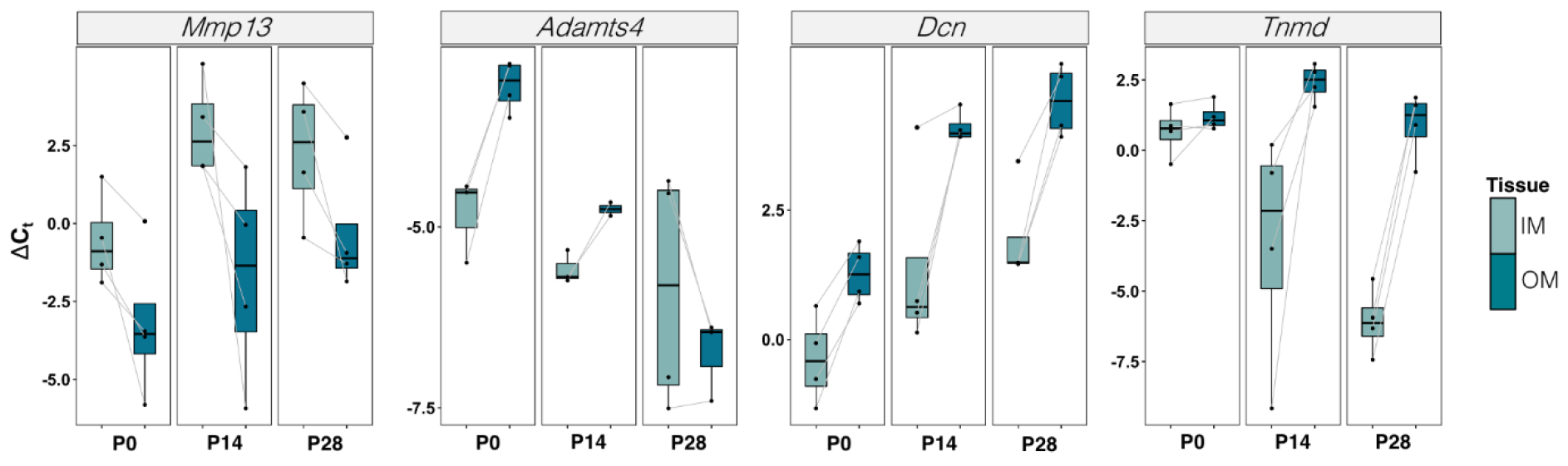
Paired gene expression (ΔC_t_) for *Mmp13*, *Adamts4*, *Dcn*, and *Tnmd* in the inner and outer meniscus (IM, OM) at P0, P14, and P28, with grey lines connecting samples from the same biologic replicate. Black line indicates mean value. Select genes show examples of gene expression patterns: heightened expression in the IM vs. OM throughout maturation (*Mmp13*), expression enrichment in the OM evident at P0 that either lessens (*Adamts4*) or persists (*Dcn*) with maturation and decrease in expression in the IM with age (*Tnmd*).

**Figure 5 Supplement 1:**
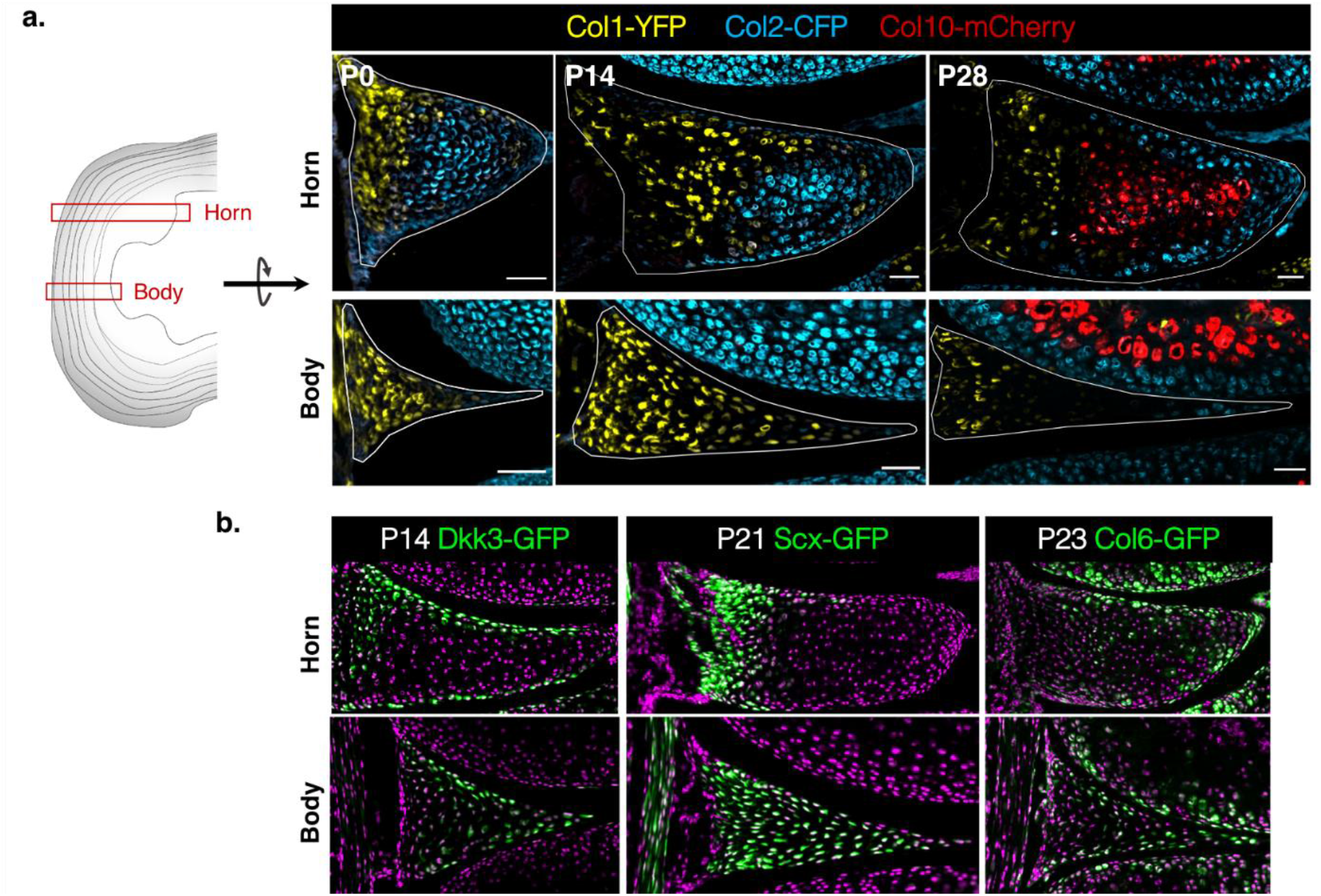
**a)** Representative coronal sections of the horn (top) and body (bottom) location within the meniscus acquired through serial sectioning of a P0, P14, and P28 meniscus. Nodules high in *Col2* reporter expression can be seen in the horn from birth, and these regions undergo hypertrophy and ossification by P28, and seen by the *Col10* reporter signal. For visualization, fluorescence intensity LUTs were not kept constant between images of different ages. Scale bar: 50 *μ*m. **b)** Representative images from the horn and body regions of 3 other transgenic mouse models: *Dkk3*-GFP (P14 sample), *Scx*-GFP (P21 sample), and *Col6a1*-GFP (P23 sample), demonstrating the potential utility of other fluorescent reporter lines in marking subpopulations within the meniscus. Cell nuclei are counterstained in magenta.

**Figure 6 Supplement 1:**
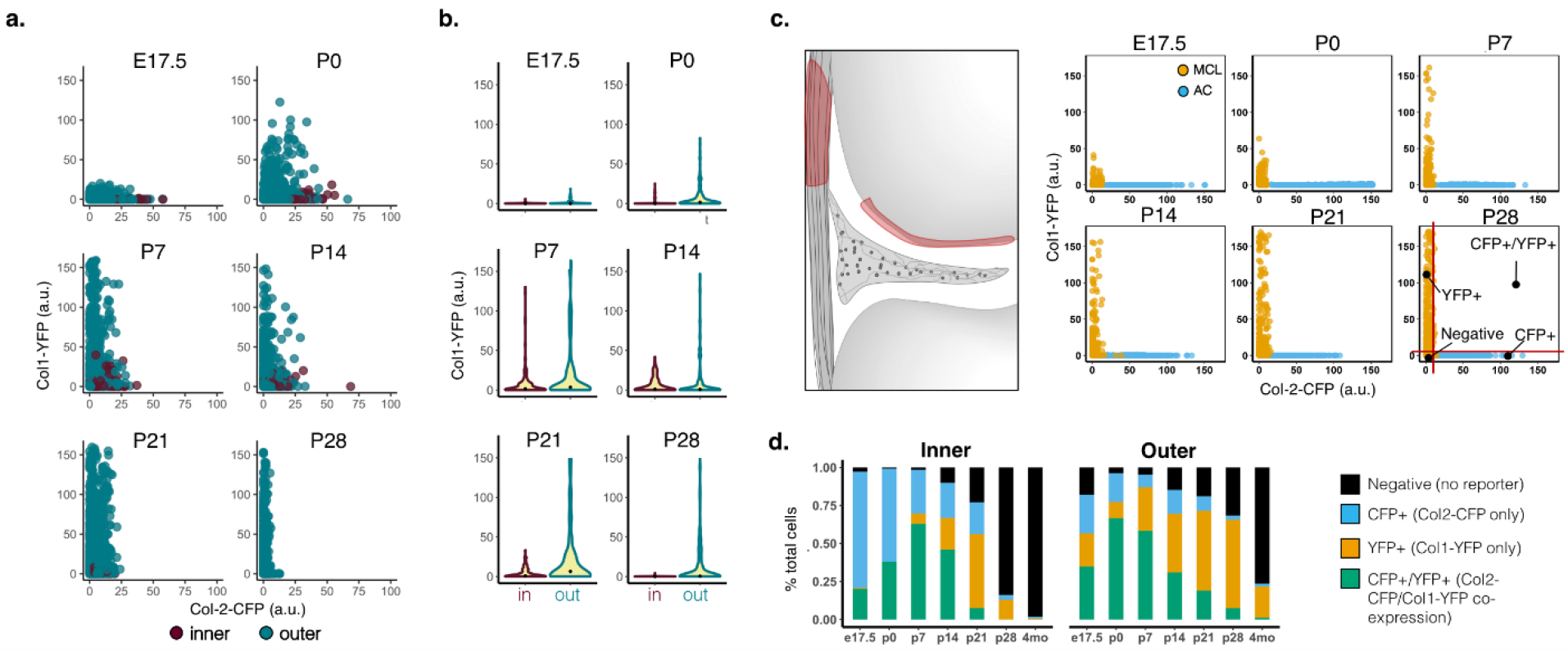
**a)** Inner and outer region Col1/Col2 reporter co-expression plots (as in Fig. 6), for all timepoints from P0-P28. **b)** Violin plots showing the distribution of measured Col1-YFP fluorescence intensities in cells of the inner and outer meniscus regions from E17.5-P28. Black dots indicate the median value. **c)** Example co-expression plots for a Col1 positive control tissue (MCL) and Col2 positive control tissue (articular cartilage). Expression of CFP in the MCL and YFP in the cartilage was assumed to be background, and these values were used to create gating criteria (red lines, P28 plot) to divide meniscus cells into the ‘Negative’, ‘CFP+’,’YFP+’, and ‘CFP+;YFP+’ populations. **d)** Percent of cells in the inner (0-0.33), middle (0.33-0.66), or outer (0.66-1) third of the meniscus that fit into one of the 4 categories in **c).** Data here is pooled for the 3 biological replicates, for ease of visualization. Differences in percentages between the inner and outer groups for each category is quantified in Fig. 6.

**Supplemental Table 1:**
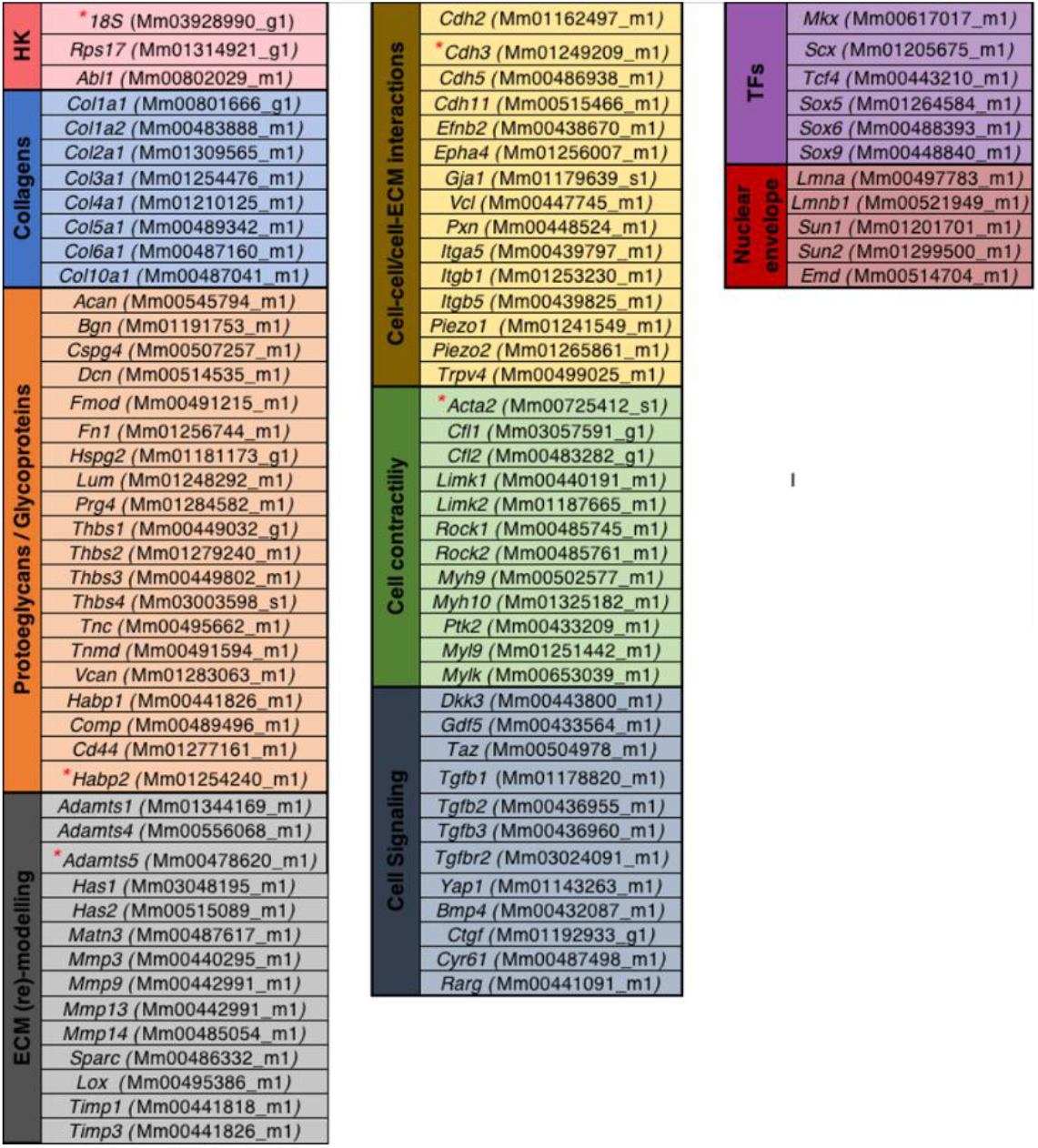
List of genes used for the Fluidigm gene expression analysis panel discussed in Fig. 3 and Fig. 4. Colored tabs indicate gene categories. HK: housekeeping, TFs: transcription factors. ThermoFisher Assay ID for each Taqman probe (primer) is listed in parentheses (see methods for more detail). Red asterisks indicate genes that were excluded based on extremely low C_t_ values (*18S*), indicating saturation of signal with pre-amplification, or genes for which more than 25% of samples contained inconclusive readouts (*Acta2, Adamts5, Cdh3, Habp2*). See methods for link to the full ClustVis analysis.

**Movie1: Z-stack of an explanted P0 meniscus visualized with forward scatter second harmonic imaging (SHG).** Circumferential aligned collagen fibers can be seen intercalating throughout the cells of the outer meniscus, while the inner meniscus region is devoid of signal— indicating an absence of organized matrix. Z-stack series shown through a 30µm depth with a step size of 0.5 µm. Imaging starts ∼15µm below the superficial tissue region. Scale bar: 25µm.

**Movie 2: SHG-visualized circumferential fibers of explanted P7 meniscus.** Forward scatter SHG through a 30µm depth (1µm step size) of a P7 explanted meniscus starting at least 15µm below the superficial surface. Close up of pockets of fibrous matrix occupied by cells shown in Fig 2 Supplement 1e. Scale bar: 25µm.

**Movie 3: SHG-visualized circumferential fibers of explanted P21 meniscus.** Forward scatter SHG through a 30µm depth (1µm step size) of a P21 explanted meniscus starting at at least 15µm below the superficial surface. Close up of pockets of fibrous matrix occupied by cells shown in Fig 2 Supplement 1e. Scale bar: 25µm.

**Movie 4**: Temporal expression of the Col-1/2/10 reporters throughout knee joint tissues largely mimics known patterns of respective endogenous collagens. Representative coronal sections of Col1-2-10 reporter mouse knees ages E17.5-4mo, imaged using equivalent illumination and fluorescence intensity scaling settings to show the relative distribution and abundance of reporter expression with age. Diagram of images tissue components in this sectioning plane shown on the left.

**Movie 5: Single-cell heterogeneity in Col-1/Col-2 reporter expression.** Whole tissue imaging of dissected P14 outer meniscus body in the transverse plane using multiphoton microscopy to capture the fibrous ECM organization (forward SHG) and heterogeneous expression profiles of the Col1-YFP and Col2-CFP reporters in the resident cells through a 30µm depth below the superficial surface. Z-step size: 1µm. Scale bar: 25µm.

